# Genetics & the Geography of Health, Behavior, and Attainment

**DOI:** 10.1101/376897

**Authors:** Daniel W Belsky, Avshalom Caspi, Louise Arseneault, David L Corcoran, Benjamin W Domingue, Kathleen Mullan Harris, Renate M Houts, Jonathan S Mill, Terrie E Moffitt, Joseph Prinz, Karen Sugden, Jasmin Wertz, Benjamin Williams, Candice L Odgers

## Abstract

People’s life chances can be predicted by their neighborhoods. This observation is driving efforts to improve lives by changing neighborhoods. Some neighborhood effects may be causal, supporting neighborhood-level interventions. Other neighborhood effects may reflect selection of families with different characteristics into different neighborhoods, supporting interventions that target families/individuals directly. To test how selection affects different neighborhood-linked problems, we linked neighborhood data with genetic, health, and social-outcome data for >7,000 European-descent UK and US young people in the E-Risk and Add Health Studies. We tested selection/concentration of genetic risks for obesity, schizophrenia, teen-pregnancy, and poor educational outcomes in high-risk neighborhoods, including genetic analysis of neighborhood mobility. Findings argue against genetic selection/concentration as an explanation for neighborhood gradients in obesity and mental-health problems, suggesting neighborhoods may be causal. In contrast, modest genetic selection/concentration was evident for teen-pregnancy and poor educational outcomes, suggesting neighborhood effects for these outcomes should be interpreted with care.

---

Young people’s life chances can be predicted by characteristics of their neighborhood ^1^. Children growing up in disadvantaged neighborhoods exhibit worse physical and mental health and suffer poorer educational and economic outcomes compared to children growing up in advantaged neighborhoods. Increasing recognition that aspects of social inequalities tend, in fact, to be geographic inequalities ^2–5^ is stimulating research and focusing policy interest on neighborhood effects and the role of place in shaping health, behavior, and social outcomes.

A challenge in interpreting neighborhood-effects research is distinguishing causal effects of neighborhood features from processes of selection in which individuals with different characteristics come to live in different neighborhoods ^6,7^. There is growing evidence that at least some neighborhood effects are causal; in a natural experiment arising from immigration policy in Sweden and in a randomized trial of a housing voucher program in the United States, people assigned to better-off neighborhoods tended to have some better health outcomes ^8,9^. Economic benefits of neighborhood interventions are less clear, but may be present for children whose neighborhoods are changed relatively early in life ^10,11^. But selection effects are also apparent. For example, in one study of hurricane survivors, those in poorer health prior to the disaster tended to relocate to higher-poverty communities in its aftermath ^12^. Selection and causation in neighborhood effects are not mutually exclusive; both can occur ^13^. Better understanding of how selection may contribute to apparent neighborhood effects is needed to guide intervention design and policy. Where selection can be ruled out as an explanation of neighborhood effects, neighborhood-level interventions could be prioritized. In instances where apparent neighborhood effects reflect selection processes, interventions delivered to individuals or families directly might prove more effective.

To evaluate the size and scope of selection effects in neighborhood research, methods are needed that quantify selection factors and that are not influenced by neighborhood conditions. The ideal approach is to compare fixed characteristics between children growing up in high-risk neighborhoods and peers growing up in better-off neighborhoods. Because neighborhoods may affect individuals as early as the very beginnings of their lives ^3,14^, traditional social-science measurements are problematic. Recent discoveries from genome-wide association studies (GWAS) provide a new opportunity to quantify selection effects at the level of the individual: polygenic scores. DNA sequence is fixed at conception and never altered by neighborhood environments. Because children inherit their DNA sequence from their parents, measures of genetic risk form a conceptual link between familial characteristics, such as parental education, that may influence selection into neighborhoods, and children’s health and social outcomes. In this article, we report proof-of-concept polygenic score analysis to quantify genetic selection into neighborhoods.

We analyzed polygenic scores and neighborhood conditions in 1,999 young people from the E-Risk Longitudinal Study, a birth cohort ascertained from a birth registry in England and Wales and followed prospectively through age 18 years. We studied phenotypes that represent substantial public health and economic burdens, have been linked with neighborhood risk in prior studies, are prevalent among 18-year-olds in England and Wales, and have been subject to large-scale genome-wide association study meta-analyses: obesity, mental health problems, teen pregnancy, and poor educational outcomes. We measured children’s genetic risk using four polygenic scores computed based on results from published GWAS of obesity, schizophrenia, age at first birth, and educational attainment ^15–18^. We measured their neighborhoods using administrative, survey, and systematic-social-observation ^19^ data collected during their childhoods. We tested for the expected associations of polygenic and neighborhood risk with E-Risk children’s development of obesity and mental-health problems, teen pregnancy, earning poor educational qualifications, and not being in education, employment, or training (NEET), as measured during home visits at age 18 years. To test for genetic selection effects, we tested for gene-environment correlations in which young people who carried elevated burdens of polygenic risk tended to have grown up in more disadvantaged neighborhoods. To test if genetic selection effects reflected the passive inheritance of genetics and neighborhood conditions from parents, we also analyzed the genetics of the children’s mothers. Finally, to test how genetics might become correlated with neighborhood conditions, we tested genetic associations with neighborhood mobility using data from 5,325 participants in the US-based National Longitudinal Study of Adolescent to Adult Health, a nationally representative longitudinal study of American adolescents followed prospectively through their late 20s/early 30s.

## RESULTS

### As anticipated by the genetics literature, E-Risk children with higher genetic risk had more social and health problems by age 18 years

We computed polygenic scores from published GWAS results for obesity, schizophrenia, age-at-first birth, and educational attainment ^15–18^ using the methods described by Dudbridge ^20^. This method proceeds as follows: First, SNPs in the E-Risk database were matched with SNPs reported in the GWAS publications. Second, for each matched SNP, a weight is calculated equal to the number of phenotype-associated alleles multiplied by the effect-size estimated in the GWAS. Finally, the average weight across all SNPs in a Study member’s genome is calculated to compute their polygenic score. Scores were transformed to have cohort-wide mean=0, standard deviation=1 for analysis.

18-year-olds with higher polygenic risk for obesity were at increased risk for obesity (RR=1.26 [1.15-1.38]); those with higher polygenic risk for schizophrenia were at increased risk for mental-health problems (RR=1.14 95%CI [1.02-1.27]); those with higher polygenic risk of young-age-at-first-birth were at increased risk for teen pregnancy (RR=1.40 95%CI [1.20-1.64]); and those with higher polygenic risk for low educational attainment were at increased risk for poor educational qualifications (RR=1.45 95%CI [1.33-1.58]) and becoming NEET (RR=1.32 [1.15-1.51]) (**Figure 1**, **Supplemental Table S1** **Panel A**).

**Figure 1.**
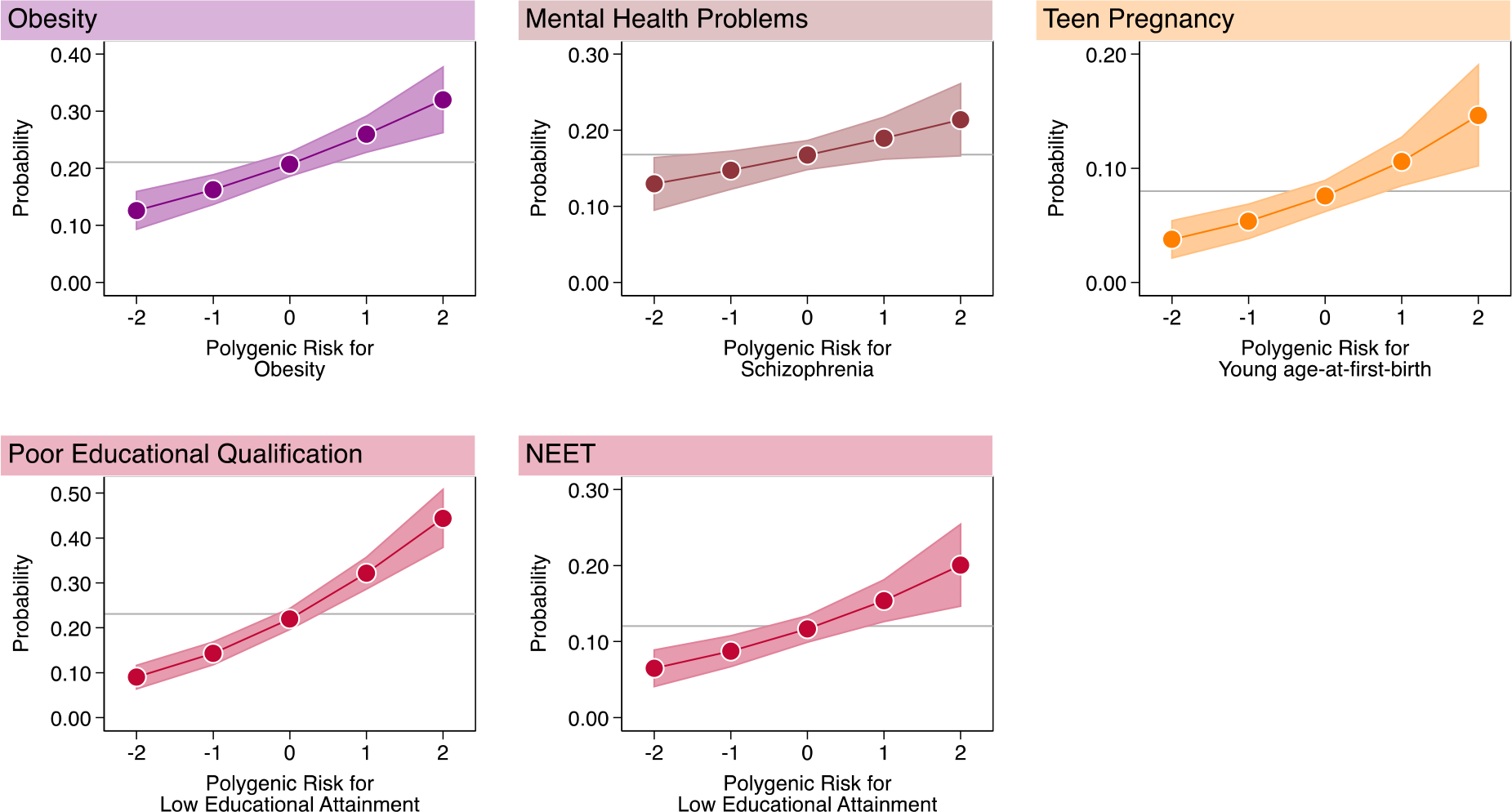
Children with higher genetic risk had more social and health problems by age 18 years. Graphs show fitted probabilities of each health and social problem across the distribution of polygenic risk. Models were adjusted for sex. Gray lines intersecting the Y-axis show the frequency of the health or social problem in E-Risk. Shaded areas around the fitted slopes show 95% Confidence intervals. Probability of obesity is graphed against polygenic risk for obesity (RR=1.26 [1.14-1.38]); probability of mental-health problems is graphed against polygenic risk for schizophrenia (RR=1.13 [1.02-1.26]); probability of teen pregnancy is graphed against polygenic risk for young age-at-first-birth 1.40 [1.19-1.64]); probabilities of poor educational qualification and NEET (Not in Education, Employment, or Training) status are graphed against polygenic risk for low educational attainment (poor educational qualification RR=1.47 [1.34-1.60]; NEET RR 1.32 [1.15-1.52]). Effect-sizes are reported for a 1-SD increase in polygenic risk.

An advantage of using genetics to test for potential selection effects is that genotypes cannot be caused by the neighborhoods where children live, ruling out reverse causation. A second advantage is that genetics may provide new information over and above what can be measured from children’s families ^21–23^. To evaluate whether the polygenic scores we studied provided new information over and above family-history risk information, we repeated our polygenic score analysis, adding covariate adjustment for family-history measures. After covariate adjustment for family history, young people’s polygenic scores remained statistically significant predictors of risk for obesity and poor educational attainment, but associations with mental health problems, teen pregnancy, and NEET status were attenuated below the alpha=0.05 threshold for statistical significance. Family history analysis is reported in **Supplemental Table S1** **Panels B and C**.

### As anticipated from the neighborhood-effects literature, children growing up in more disadvantaged neighborhoods were at increased risk for social and health problems by age 18 years

Because there is no single standard to quantify neighborhood risks ^6,24^, we used two different approaches to measure E-Risk children’s exposure to neighborhood disadvantage.

The first approach characterized the neighborhoods as they are seen by businesses and the public sector, using a consumer-classification system called ACORN (“A Classification of Residential Neighborhoods”). We computed the average ACORN classification across children’s home addresses when they were aged 5, 7, 10, and 12 years. According to ACORN, 22% of E-Risk cohort children grew up in “Wealthy Achiever” neighborhoods, 33% grew up in “Urban Prosperity/ Comfortably Off” neighborhoods, 19% grew up in “Moderate Means” neighborhoods, and 26% grew up in “Hard Pressed” neighborhoods. This distribution matched overall distributions for the United Kingdom. As an example, ACORN distributions for E-Risk families at the time of the age-12 assessment are compared to the national distribution in **Figure 2**, **Panel A**.

**Figure 2.**
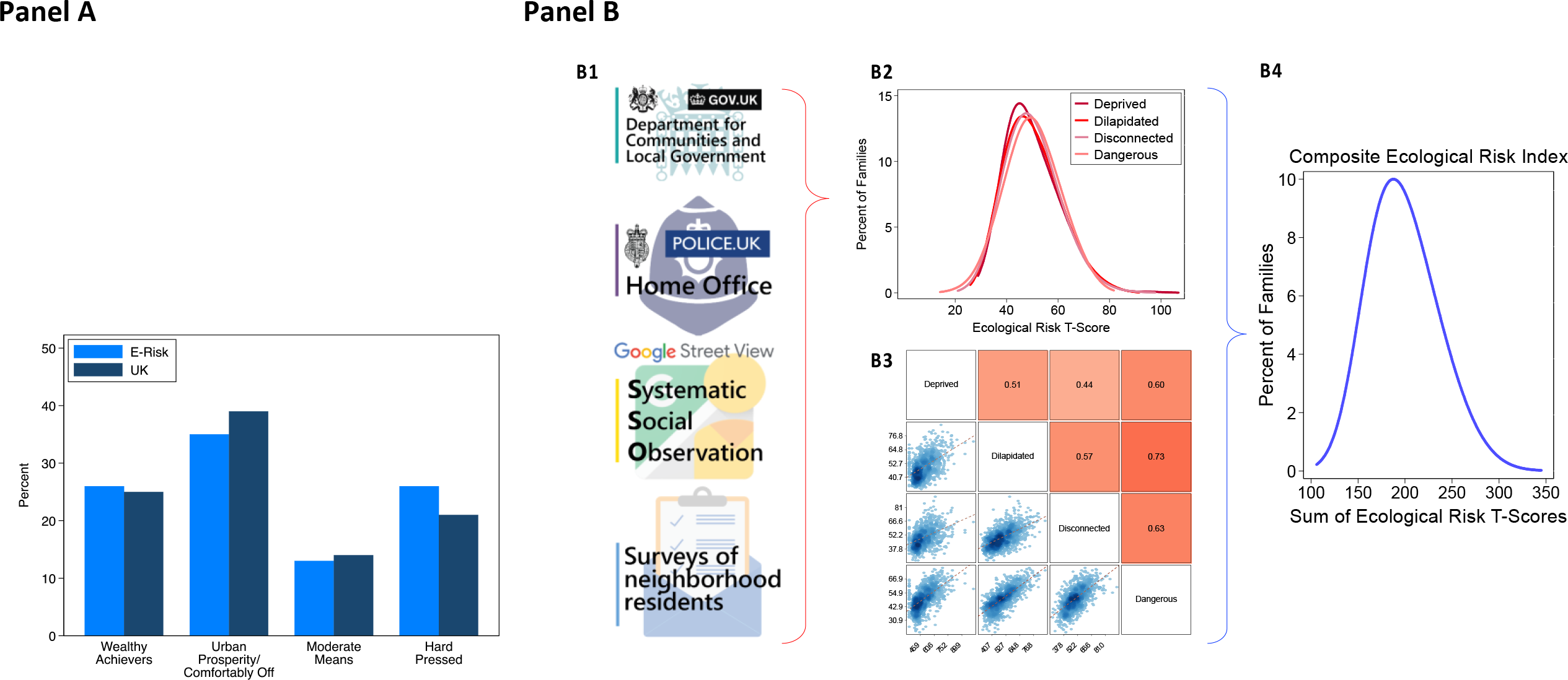
Quantification of E-Risk families’ neighborhood disadvantage using ACORN and a composite Ecological-Risk Index. **Panel A** of the Figure shows distributions of ACORN (“A Classification of Residential Neighborhoods”) classifications for E-Risk families at the time of the age-12 interview (light blue bars) and the corresponding distribution for the United Kingdom obtained from http://doc.ukdataservice.ac.uk/doc/6069/mrdoc/pdf/6069_acorn_userguide.pdf (dark blue bars). **Panel B** of the Figure contains 4 cells. Cell B1 depicts the four sources of data used for ecological-risk assessment: from top to bottom, these are geodemographic data from local governments, official crime data, Google Streetview Systematic Social Observation (SSO), and resident surveys. Cell B2 shows distributions of four ecological-risk measures derived from these data: economic deprivation, physical dilapidation, social disconnectedness, and danger. Values of the ecological-risk measures are expressed as T scores (M=50, SD=10). Cell B3 shows a matrix of the ecological-risk measures. Matrix cells below and to the left of measures show scatterplots of their association. Matrix cells above and to the right of measures show their correlation expressed as Pearson’s r. illustrating their correlation with one another (Pearson r = 0.4-0.7). Cell B4 shows the distribution of the composite Ecological-Risk Index calculated as the sum of ecological-risk measure T scores (M=198, SD=33).

The second approach characterized neighborhoods as they are seen by social scientists and public health researchers. Ecological risk measures were constructed from (a) geodemographic data from local governments, (b) official crime data accessed as part of an open data sharing effort about crime and policing in England and Wales, (c) Google Streetview Virtual Systematic Social Observation ^19^, and (d) data from surveys of neighborhood residents. We used these data to score neighborhoods on their economic deprivation, physical dilapidation, social disconnection, and dangerousness. We standardized scores to have M=50, SD=10 (“T” scores). We summed these four ecological-risk measures to compute one composite Ecological-Risk Index (M=198, SD=33) (**Figure 2 Panel B**). This composite Ecological-Risk Index was correlated with ACORN classifications, r=0.65 (**Supplemental Figure 1**; Correlations among all neighborhood measures are reported in **Supplemental Table 2**).

18-year-olds who grew up in neighborhoods with more disadvantaged ACORN classifications or with higher scores on the Ecological-Risk Index were at increased risk for obesity, mental-health problems, teen pregnancy, poor educational qualifications, and NEET status (obesity ACORN RR=1.22 [1.11-1.34], Ecological-Risk Index RR=1.15 [1.03-1.29]; mental health problems ACORN RR=1.21 [1.08-1.35], Ecological-Risk Index RR=1.30 [1.14-1.47]; teen pregnancy ACORN RR=1.64 [1.38-1.94], Ecological-Risk Index RR=1.55 [1.30-1.85]; poor educational qualifications ACORN RR=1.59 [1.44-1.76], Ecological-Risk Index RR=1.47 [1.33-1.62]; NEET ACORN RR=1.58 [1.37-1.84], Ecological-Risk Index RR=1.59 [1.36-1.85]). **Figure 3** plots risk for each health and social problem by childhood neighborhood disadvantage. Results for all neighborhood measures are reported in **Supplemental Table 3**.

**Figure 3.**
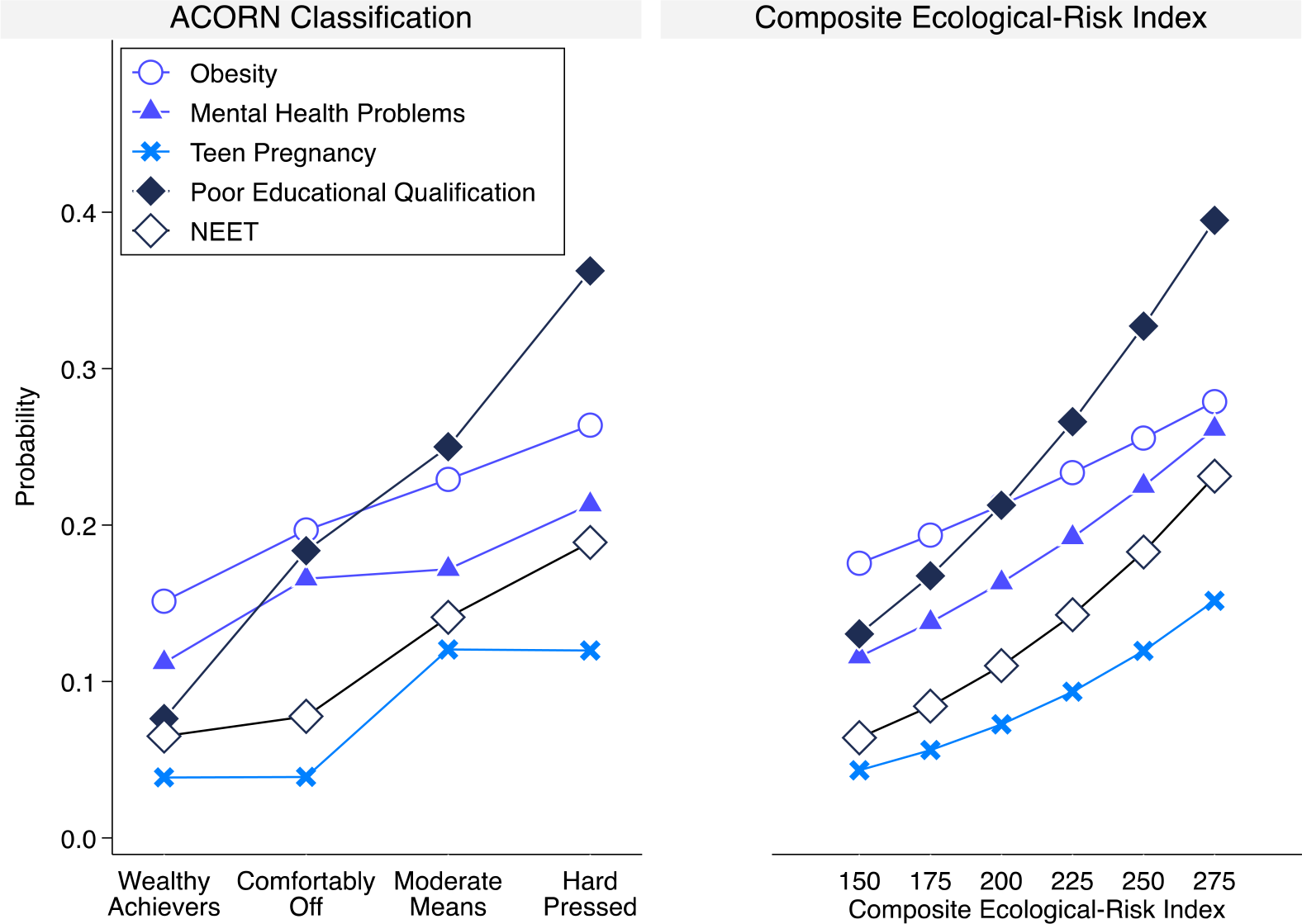
Children growing up in more disadvantaged neighborhoods were at increased risk for social and health problems by age 18 years. Graphs show the neighborhood risk gradient for each health and social problem. The y-axis shows the probability of having a given problem at varying levels of neighborhood risk. The left-side graph plots probabilities by ACORN Classification. The right-side graph plots predicted probabilities for a series of values of the composite Ecological-Risk Index. (The average neighborhood is expected to have a risk score of 200. A composite Ecological-Risk Index of 150 is about one and a half standard deviations below the mean for E-Risk family neighborhoods.) Effect-sizes in terms of relative risk associated with a 1-cateogry increase in disadvantage classified by ACORN / 1-standard deviation increase in composite Ecological-Risk Index were: obesity (RR=1.20 [1.10-1.31] / 1.15 [1.03-1.28]); mental health problems (RR=1.19 [1.08-1.31]/ 1.27 [1.12-1.44]); having a teen pregnancy (RR=1.56 [1.34-1.83]/ 1.52 [1.27-1.81]); poor educational qualifications (RR=1.53 [1.40-1.67]/ 1.48 [1.34-1.63]) and NEET (RR=1.52 [1.33-1.74]/ 1.57 [1.35-1.83]) (**Supplemental Table 1**).

### We found little evidence that genetic selection/concentration explained neighborhood risk for obesity or mental health problems

Although children’s genetic risk and their neighborhood disadvantage separately predicted their increased risk of obesity and mental health problems, polygenic risks for obesity and schizophrenia were not consistently related to neighborhood disadvantage. **Panels A and B of Figure 4** show this result graphically. Whereas the blue slopes document positive associations between neighborhood disadvantage (x-axes) and risk for obesity (left-side y-axes of Panel A) and mental health problems (left-side y-axes of Panel-B), the red slopes reveal null associations between neighborhood disadvantage (x-axes) and polygenic risk for obesity (right-side y-axes of Panel A) and null or weak associations between neighborhood disadvantage and polygenic risk for schizophrenia (right-side y-axes of Panel B). In the E-Risk cohort, children raised in disadvantaged neighborhoods more often became obese by age 18; but we found no evidence for concentration of children with high polygenic risk in disadvantaged neighborhoods (ACORN r=−0.02, p=0.561; Ecological-Risk Index r=0.00, p=0.985). Results were similar for analysis of genetic risk for schizophrenia, although the association between children’s polygenic scores and their neighborhood Ecological-Risk Index was statistically significant at the α=0.05 level (ACORN r=0.04, p=0.178; Ecological-Risk Index r=0.07, p=0.049). Results for all neighborhood measures are reported in **Supplemental Table 4**. These findings argue against neighborhood selection/composition as a source of neighborhood gradients in obesity and mental health problems and encourage more research to unravel the possible causal effects of neighborhood conditions on physical and mental health.

**Figure 4.**
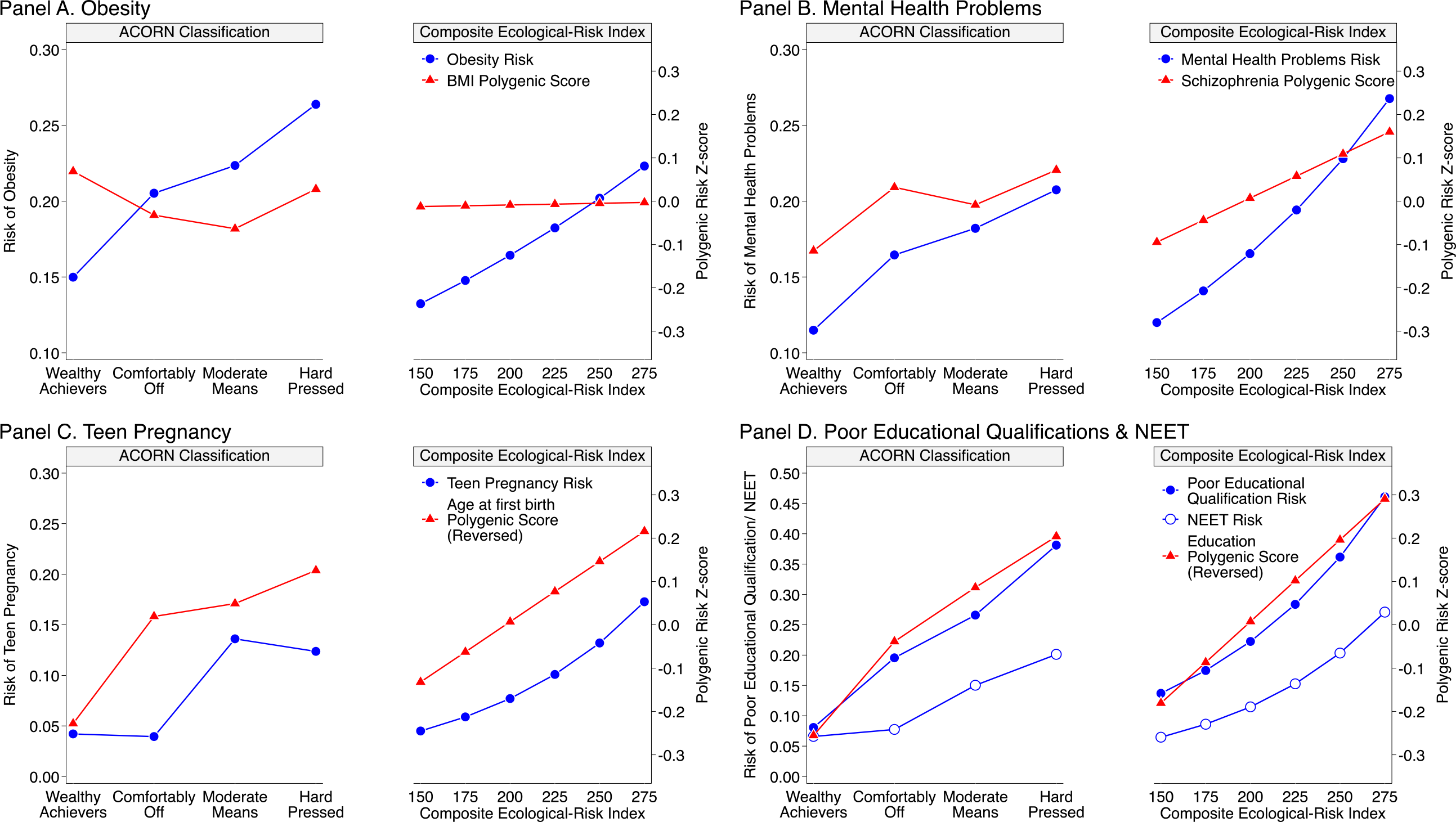
Neighborhood gradients in phenotypic and genetic risk for obesity, mental health problems, teen pregnancy, poor educational qualifications, and NEET status. Figures graph E-Risk young adults’ phenotypic risk (blue slopes, left-side y-axes showing risk expressed as a predicted probability) and genetic risk (red slopes, right-side y-axes showing polygenic risk expressed on Z scale) across the distribution of neighborhood disadvantage (x-axes). The left figure panes graph results for ACORN classification of neighborhood disadvantage. The right figure panes graph results for ecological-risk score quantification of neighborhood disadvantage. **Panel A** graphs risk of obesity (blue) and body-mass-index (BMI) polygenic score, red). The panel shows that E-Risk participants growing up in more disadvantaged neighborhoods more often became obese, but did not differ from peers in their polygenic risk for obesity. **Panel B** graphs risk of mental health problems (blue) and schizophrenia polygenic score (red). The panel shows that E-Risk participants growing up in more disadvantaged neighborhoods more often developed mental health problems and were at higher polygenic risk for schizophrenia, although the genetic association was statistically significant only for the ecological-risk score measure of neighborhood disadvantage. **Panel C** graphs risk of teen pregnancy (blue) and age-at-first-birth polygenic score (red). Age-at-first-birth polygenic score values are reversed for the graph so that higher values correspond to genetic prediction of younger age at first birth. The panel shows that E-Risk participants growing up in more disadvantaged neighborhoods more often had teen pregnancies and had higher polygenic risk for early first birth. **Panel D** graphs risk of poor educational qualifications (blue dots) and NEET status (blue circles), and educational attainment polygenic score (red). Education polygenic score values are reversed for the graph so that higher values correspond to genetic prediction of lower educational attainment. The panel shows that E-Risk participants growing up in more disadvantaged neighborhoods more often struggled with education and employment and tended to have higher polygenic risk for low educational attainment.

### We found evidence of genetic selection/concentration in disadvantaged neighborhoods of children at high polygenic risk of teen pregnancy, poor educational attainment, and NEET

We tested if children at higher polygenic risk for young age-at-first-birth and poor educational attainment tended to grow up in more disadvantaged neighborhoods. They did, as measured by both ACORN classification and the composite Ecological-Risk Index. **Panels C and D of Figure 4** show this result graphically. In Panel C, the blue slopes document the positive association between neighborhood disadvantage (x-axis) and risk of teen pregnancy (left-side y-axis); the red slope illustrates the positive association between neighborhood disadvantage and polygenic risk for young age-at-first-birth (reversed values of the age-at-first-birth polygenic score, right-side y-axis; ACORN r=0.12, p<0.001; Ecological-Risk Index r=0.11, p=0.003). In Panel D, the blue slopes document the positive association between neighborhood disadvantage (x-axis) and poor educational attainment and NEET status; the red slopes illustrate the positive association between neighborhood disadvantage and polygenic risk for low educational attainment (reversed values of the education polygenic score, right-side y-axis; ACORN r=0.19, p<0.001; Ecological-Risk Index r=0.15, p<0.001). Results for all neighborhood measures are reported in **Supplemental Table 4**. These findings suggest that neighborhood selection/composition may be relevant to neighborhood-teen pregnancy and neighborhood-achievement gradients and encourage research to understand selection processes.

### Children inherited genetic and neighborhood risks from their parents

E-Risk children were aged 5-12 years during the period when neighborhood data were collected. It is unlikely that they actively selected themselves into different types of neighborhoods. Instead, a hypothesis for why children’s polygenic and neighborhood risks are correlated is that both risks are inherited from their parents. According to this hypothesis, genetics influence parents’ characteristics and behaviors, which in turn affect where they live. Children subsequently inherit their parents’ genetics and their neighborhoods. As an initial test of this hypothesis, we analyzed genetic data that we collected in the E-Risk study from children’s mothers (N=840 with children included in analysis). (E-Risk did not collect fathers’ DNA.) As expected, polygenic scores were correlated between E-Risk participants and their mothers (r=0.49-0.52). We first tested if mothers’ polygenic scores were associated with neighborhood disadvantage. Parallel to results from analysis of children’s genetics, mothers’ polygenic risk for obesity and schizophrenia were not associated with their neighborhood disadvantage (r=−0.01-0.03, p>0.3). Also consistent with analysis of children, mother’s age-at-first-birth and educational-attainment polygenic scores were associated with neighborhood disadvantage (effect-sizes r=0.14-0.20, p<0.001 for all; **Supplemental Table 5**). Next, we repeated analysis of association between children’s polygenic scores and neighborhood disadvantage, this time including a covariate for the mother’s polygenic score. Consistent with the hypothesis that children’s polygenic and neighborhood risks are correlated because both risks are inherited from their parents, covariate adjustment for mothers’ polygenic scores reduced magnitudes of associations between children’s polygenic scores and their neighborhood disadvantage by more than half (**Supplemental Table 6**, **Supplemental Figure 2**).

### Polygenic risk for teen pregnancy and low educational attainment predicted downward neighborhood mobility among participants in the US National Longitudinal Study of Adolescent to Adult Health

If children’s genetic and neighborhood risks are correlated because they inherit both risks from their parents, the next question is how parents’ genetics come to be correlated with neighborhood risks. A hypothesis is that parents’ genetics influence their characteristics and behavior in ways that affect where they are able to live. To test this hypothesis, data are needed that observe the neighborhood mobility process in which people leave the homes where they grew up and selectively end up in new neighborhoods. Because the E-Risk study began collecting information on children’s mothers only after the children were born, data were not collected on the mothers’ own childhood neighborhoods. Therefore, to test how polygenic risks might influence patterns of neighborhood mobility, we turned to a second dataset, the US National Longitudinal Study of Adolescent to Adult Health (Add Health). Add Health first surveyed participants when they were secondary-school students living with their parents. Add Health has since followed participants into their late 20s and early 30s ^25^, when most were living in new neighborhoods (N=5,325 with genetic and neighborhood data; 86% lived >1km from the address where they first surveyed). We used the Add Health genetic database and neighborhood measures derived from US Census data to test if polygenic risk for obesity, schizophrenia, teen-pregnancy, and low educational attainment predicted downward neighborhood mobility, i.e. young adults coming to live in more disadvantaged neighborhoods relative to the one where they lived with their parents.

Add Health participants’ polygenic risks for obesity and schizophrenia showed weak or null associations with neighborhood disadvantage when they were first surveyed in their parents’ homes as secondary school students (polygenic risk for obesity r=0.03, p=0.028; polygenic risk for schizophrenia r=−0.01, p=0.675) and when they were followed-up in their 20s and 30s (polygenic risk for obesity r=0.04, p=0.023; polygenic risk for schizophrenia r=−0.03, p=0.063). Findings were similar in neighborhood mobility analysis (polygenic risk for obesity r=0.03, p=0.036; polygenic risk for schizophrenia r=−0.02, p=0.072. These findings bolster conclusions from E-Risk analysis that genetic selection/concentration is likely to be a trivial factor in neighborhood gradients in obesity and mental health problems, although the obesity polygenic score association with neighborhood risk was statistically significant in Add Health, and therefore not a full replication of findings in E-Risk.

In contrast, Add Health participants with higher polygenic risk for teen pregnancy and low educational attainment tended to have grown up in more disadvantaged neighborhoods (r=0.07 for the age-at-first-birth polygenic score and r=0.17 for the educational-attainment polygenic score; p<0.001 for both) and to live in more disadvantaged neighborhoods when they were followed-up in their 20s and 30s (adult neighborhood r=0.09 for the age-at-first-birth polygenic score and r=0.13 for the educational attainment polygenic score; p<0.001 for both). In neighborhood mobility analysis, participants with higher polygenic risk for teen pregnancy and low educational attainment tended to move to more disadvantaged neighborhoods relative to the neighborhoods where they lived with their parents when they were first surveyed (downward mobility r=0.06 for the age-at-first-birth polygenic score and r=0.07 for the educational-attainment polygenic score; p<0.001 for both; **Figure 6**). These finding bolster conclusions from E-Risk analysis that genetic selection/concentration may contribute to neighborhood gradients in teen pregnancy and poor educational outcomes, although this contribution may be small.

### Summary

Sociogenomic analyses testing the concentration of polygenic risks for health, behavior, and social problems in children growing up in disadvantaged neighborhoods yielded three findings: We found little consistent evidence for the concentration of polygenic risk for obesity or polygenic risk for mental health problems in children growing up in disadvantaged neighborhoods. In contrast, we found consistent evidence for the concentration of polygenic risks for teen pregnancy and low achievement. Concentration of polygenic risks was mostly explained by children’s inheritance of both neighborhood and polygenic risks from their parents. Selective mobility may contribute to concentrations of risks. In neighborhood mobility analysis that followed young people living with their parents during adolescence to where they lived as adults nearly two decades later, participants with higher polygenic risk for teen pregnancy and low achievement exhibited downward neighborhood mobility, moving to more disadvantaged neighborhoods across follow-up.

## DISCUSSION

Large investments are being made in neighborhood-level policies and programs intended to improve the health and wellbeing of residents. These investments are based on exciting new findings demonstrating causal long-term impacts of local neighborhoods on health and possibly economic outcomes for children moving out of poverty ^9–11^. The promise of place-based intervention efforts is that they can improve, at scale, the lives of residents and, for children, break the intergenerational transmission of poverty and lack of opportunity. At the same time, genome-wide association studies (GWAS) are revealing genetic predictors of the health outcomes, behaviors, and attainments that place-based interventions seek to modify. We carried out a study of genetic selection into neighborhoods to test how genes and geography combine.

### We did not find consistent evidence of genetic selection effects into neighborhoods for obesity and mental health problems

To our knowledge, this is the first study testing gene-neighborhood correlations with GWAS discoveries for obesity. A previous Swedish study detected evidence of gene-neighborhood correlation between the schizophrenia polygenic score and a commercial database measure of neighborhood deprivation (r=0.04, N^~^7,000) ^26^. The magnitude of the association was about the same as we observed for the E-Risk twins (r=0.04) and their mothers (r=0.03) using the commercial-database ACORN measure. However, this association was not replicated in the Add Health Study, where the association was not statistically significant and was in the opposite direction (r=−.01 to −.03). It is possible that selective non-participation in research related to genetic liability for schizophrenia could limit ability to detect these associations in some datasets ^27,28^. However, selective participation related to genetics has been documented for other polygenic scores, including the educational attainment polygenic score ^29^. Our results nevertheless document consistent evidence (across measures and samples) of gene-neighborhood correlation for GWAS discoveries for age-at-first-birth and educational attainment, and less so for GWAS discoveries for obesity and schizophrenia.

### We found consistent evidence of genetic selection effects into neighborhoods for teen pregnancy, poor educational qualifications, and NEET status

These findings are consistent with recent findings in sociology about how neighborhood residents come to be both physically and economically “stuck in place” across generations ^30,31^. Teen pregnancy and poor outcomes in education and the workplace can trap parents and their children in disadvantaged neighborhoods, causing a clustering of individual-level and neighborhood-level risks. This has led to calls for multi-generational and multi-level intervention efforts to break the cycle of disadvantage. While our findings show that selection is at work for these key outcomes, the effects documented are unlikely to be large enough to fully account for neighborhood gradients. Consistent evidence for both selection (from us) and social causation (in the larger literature) means that policies and interventions will need to target resources at both people and place to be effective.

Our findings make three contributions. First, they make a methodological and conceptual contribution by integrating genetics and social science in the rapidly developing field of social geography. We know that the places where children grow up are associated with whether they thrive. The challenge in neighborhood research is to sort out selection from causation. Here, we take a fresh look at this classic problem using new information from genomics research. DNA-sequence differences between people index differences in liability to health and social outcomes, and DNA cannot be influenced by neighborhoods. As the price of generating genetic data continues to fall, measurements of these DNA differences can provide tools to advance social science research into effects of place.

Second, findings shed light on how genetics and environments combine to influence children’s development. Genetics contribute to the effects of place by influencing where people choose to live, are forced to live, or otherwise end up living. For E-risk and Add Health young people, some genetic risks were patterned across neighborhoods, presumably reflecting the children’s inheritance of genetics that influenced where their parents were able to live. This patterning was apparent for genetics linked to teen pregnancy and poor education, but not with genetics linked to mental health problems or obesity. One interpretation is that teen pregnancy and poor education are more proximate causes of economic circumstances that determine where one can live as compared to, for example, obesity. Consistent with this interpretation, Add Health young people who carried higher levels of polygenic risk for teen pregnancy and poor educational outcomes showed patterns of downward neighborhood mobility, tending to move in young adulthood to worse-off neighborhoods relative to the ones where they grew up. In contrast, Add Health participants’ polygenic risk for obesity and schizophrenia showed trivial or null associations with their neighborhood mobility. Findings document that even though risk for highly heritable health problems such as obesity and schizophrenia may be patterned across neighborhoods, genetic risks for these conditions may not be. More broadly, findings highlight that a phenotype being heritable does not imply that social risk factors are necessarily genetically confounded.

Third, findings provide evidence that many children are growing up subject to correlated genetic and place-related risks, particularly for teen pregnancy and attainment failure. The genetic selection effects we observed are too small to account entirely for neighborhood effects, but genetic and neighborhood risks may act in combination. Neighborhood interventions can thus be conceptualized, in part, as breaking up gene-environment correlations, lending urgency to the development of effective place-based interventions. To this end, genetically-informed designs may offer opportunities to advance intervention research. For example, comparative studies could test if correlations between genetic and neighborhood risks vary across cities governed by different urban planning strategies. Intervention studies could also actively incorporate genetic information: Trials of neighborhood interventions can improve precision of their treatment-effect estimates by including polygenic score measurements as control variables to account for unmeasured differences between participants ^32^.

We acknowledge limitations. Foremost, our measures of genetic risk are imprecise. They explain only a fraction of the genetic variance in risk estimated from family-based genetic models; the polygenic scores for educational attainment explains >10% of phenotypic variance, polygenic scores for body-mass index and schizophrenia explain 6-7% of phenotypic variance, and the age-at-first-birth polygenic score explains one percent of phenotypic variance ^15–18^, whereas heritabilities of these traits and behaviors estimated in family-based studies tend to be much higher ^33^. As a consequence, our estimates of gene-neighborhood correlations should be considered lower-bound estimates. Second, a related limitation is that the different polygenic scores had different amounts of power to detect associations with neighborhood risk, with the education polygenic score having more power than the others. Nevertheless, we had more power for body-mass index and schizophrenia polygenic score analysis than we did for age-at-first-birth polygenic score analysis, and yet genetic associations with neighborhood risk were much larger for the age-at-first-birth polygenic score than for the body-mass index and schizophrenia polygenic scores. This pattern held in both the E-Risk and Add Health studies. Third, the magnitudes of observed gene-neighborhood correlations in our study were small. For example, the strongest gene-neighborhood correlations we observed were for the educational attainment polygenic score (r^~^0.17). Based on our analysis, neighborhood differences in this polygenic score between the highest and lowest risk neighborhoods could account for, at most, only about 15% of the observed differences in poor educational qualifications and about 10% of the observed differences in NEET status between these neighborhoods (**Supplemental Methods**). As GWAS sample sizes continue to grow, more precise measurements will become available ^34^. More predictive polygenic scores could potentially strengthen measured gene-neighborhood correlations and explain increasing fractions of neighborhood gradients in health and social outcomes.

An additional limitation is that E-Risk data come from a single birth cohort in a single country, and thus reflect a relatively specific geographic and historical context. Findings that polygenic risk of teen pregnancy and low educational attainment were correlated with neighborhood disadvantage did replicate in the US-based Add Health Study. Add Health neighborhood risk was measured from tract-level US Census data describing broad social and economic conditions and is thus less geographically precise than the small-area ACORN and Ecological-Risk Assessment data analyzed in E-Risk. Therefore, Add Health analysis is not a direct replication of our E-Risk findings. Instead, the consistent results across two studies of different populations measured using different methods argues for the overall robustness of our findings.

Finally, our analysis in both the E-Risk and Add Health studies was limited to European-descent individuals. This restriction was necessary to match the ancestry of our analytic sample with the ancestry of the samples studied in the GWAS used to calculate polygenic scores, which is the recommended approach ^35^. As polygenic scores are developed for populations of non-European ancestry, replication in these populations should be a priority.

The observation of gene-neighborhood correlations does not suggest that residents in disadvantaged neighborhoods will not benefit from neighborhood-level interventions. It simply means policy-makers should not over-interpret neighborhood effects in purely causal terms. For example, people observed to live in a friendly suburb, remote ranch, quaint village, and luxury high-rise are not found in those neighborhoods randomly by accident; people end up in such locations selectively. But regardless of location they all respond to incentives and opportunities. More precise quantifications of selection processes influencing where people live can help inform policies and programs to craft incentives and opportunities that promote healthy development for everyone.

## METHODS

### Environmental Risk Longitudinal Study (E-Risk)

#### Sample

Participants were members of the Environmental Risk (E-Risk) Longitudinal Twin Study, which tracks the development of a birth cohort of 2,232 British children. The sample was drawn from a larger birth register of twins born in England and Wales in 1994–1995 ^36^. Full details about the sample are reported elsewhere ^37^. Briefly, the E-Risk sample was constructed in 1999-2000, when 1,116 families (93% of those eligible) with same-sex 5-year-old twins participated in home-visit assessments. The sample includes 56% monozygotic and 44% dizygotic twin pairs; sex is evenly distributed within zygosity (49% male). Families were recruited to represent the UK population of families with newborns in the 1990s, on the basis of residential location throughout England and Wales, and mother’s age. Teenaged mothers with twins were over-selected to replace high-risk families who were selectively lost to the register through nonresponse. Older mothers having twins via assisted reproduction were under-selected to avoid an excess of well-educated older mothers. These strategies ensured that the study sample represents the full range of socioeconomic conditions in Great Britain ^19^.

Follow-up home visits were conducted when the children were aged 7 (98% participation), 10 (96% participation), 12 (96% participation), and, in 2012–2014, 18 years (93% participation). There were no differences between those who did and did not take part at age 18 in terms of socioeconomic status (SES) assessed when the cohort was initially defined (χ^2^ = 0.86, *p* = .65), age- 5 IQ scores (*t* = 0.98, *p* = .33), or age- 5 internalizing or externalizing behavior problems (*t* = 0.40, *p* = .69 and *t* = 0.41, *p* = .68, respectively). Home visits at ages 5, 7, 10, and 12 years included assessments with participants as well as their mother; the home visit at age 18 included interviews only with the twin participants. All interviews at the age-18 assessment were conducted after the 18th birthday. Each twin participant was assessed by a different interviewer. The joint Research and Development Office of South London and Maudsley and the Institute of Psychiatry Research Ethics Committee approved each phase of the study. Parents gave informed consent and twins gave assent between ages 5 and 12 years; twins gave informed consent at age 18 years.

#### Genetic Data

We used Illumina HumanOmni Express 12 BeadChip arrays (Version 1.1; Illumina, Hayward, CA) to assay common single-nucleotide polymorphism (SNP) variation in the genomes of cohort members. We imputed additional SNPs using the IMPUTE2 software (Version 2.3.1; https://mathgen.stats.ox.ac.uk/impute/impute_v2.html; ^38^) and the 1000 Genomes Phase 3 reference panel ^39^. Imputation was conducted on autosomal SNPs appearing in dbSNP (Version 140; http://www.ncbi.nlm.nih.gov/SNP/; ^40^) that were “called” in more than 98% of the samples. Invariant SNPs were excluded. Pre-phasing and imputation were conducted using a 50-million-base-pair sliding window. We analyzed SNPs in Hardy-Weinberg equilibrium (*p* > .01). The resulting genotype databases included genotyped SNPs and SNPs imputed with 90% probability of a specific genotype among European-descent E-Risk Study members (N=1,999 children in 1,011 families). The same procedure was used to construct the genetic database for Study members’ mothers. Genetic data were available for N=840 mothers of the Study members in our genetic analysis sample.

#### Polygenic Scoring

We computed polygenic scores for obesity (body-mass index), schizophrenia, age-at-first birth, and educational attainment from published genome-wide association study (GWAS) results ^15–18^. We computed these polygenic scores because the GWAS on which they are based are among the largest and most comprehensive available and their target phenotypes are established as having strong geographic gradients in risk. For example, in the case of schizophrenia, there is a long-running debate about hypotheses of social causation, in which ecological risks contribute to schizophrenia pathogenesis, and social drift, in which genetic liability to schizophrenia causes downward social mobility^41,42^.

Polygenic scoring was conducted following the method described by Dudbridge ^20^ using the PRSice software ^43^. Briefly, SNPs reported in GWAS results were matched with SNPs in the E-Risk database. For each SNP, the count of phenotype-associated alleles (i.e. alleles associated with higher body-mass index, increased risk of schizophrenia, younger age at first birth, or less educational attainment, depending on the score being calculated) was weighted according to the effect estimated in the GWAS. Weighted counts were summed across SNPs to compute polygenic scores. We used all matched SNPs to compute polygenic scores irrespective of nominal significance in the GWAS.

Polygenic score analysis may be biased by population stratification, the nonrandom patterning of allele frequencies across ancestry groups ^35,44^. To address residual population stratification within the European-descent members of the E-Risk sample, we conducted principal components analysis ^45^. We computed principal components from the genome-wide SNP data with the PLINK software ^46^ using the command ‘pca’. One member of each twin pair was selected at random from each family for this analysis. SNP-loadings for principal components were applied to co-twin genetic data to compute principal component values for the full sample. We residualized polygenic scores for the first ten principal components estimated from the genome-wide SNP data ^47^ and standardized residuals to have mean=0, SD=1 for analysis.

#### Neighborhood Disadvantage

We characterized the neighborhoods in which E-Risk participants grew up using two approaches. The first approach characterized the neighborhoods as they are seen by businesses and the public sector, using a consumer-classification system called ACORN (“A Classification of Residential Neighborhoods”). The second approach characterized neighborhoods as they are seen by social scientists and public health researchers, using ecological-risk assessment methods.

##### Neighborhood Disadvantage Measured by Consumer Classification

We used a geodemographic classification system, ACORN, developed as a tool for businesses interested in market segmentation by CACI (CACI, UK, http://www.caci.co.uk/). This is a proprietary algorithm that is sold to businesses, but which CACI made available to our research group. ACORN classifications were derived from analysis of census and consumer research databases. ACORN classifies neighborhoods, in order of least disadvantaged to most disadvantaged as “Wealthy Achievers”, “Urban Prosperity”, “Comfortably Off”, “Moderate Means”, or “Hard Pressed”. For analysis, we combined neighborhoods classified in the “Urban Prosperity” and “Comfortably Off” categories because very few children lived in “Urban Prosperity” neighborhoods (Nationally, fewer children live in neighborhoods characterized by “Urban Prosperity”). We obtained ACORN classifications for the Output Areas in which E-Risk families’ lived ^19^. Output Areas are the smallest unit at which UK Census data are provided and they reflect relatively small geospatial units of about 100-125 households. Households were classified based on street address at the time of the age-5, age-7, age-10, and age-12 in-home visits. We assigned children the average neighborhood classification across these four measurements. ACORN classifications were available for N=1,993 children in 1,008 families in the genetic sample.

##### Neighborhood Disadvantage Measured by Ecological-Risk Assessment

Ecological risk assessment was conducted by combining information from four independent sources of data: Geodemographic data from local governments, official crime data from the UK police, Google Streetview-based Systematic Social Observation (SSO), and surveys of neighborhood residents conducted by the E-Risk investigators when the E-Risk children were aged 13-14. These data sources are described in detail in the **Supplementary Methods**.

We used these four data sources to measure neighborhood ecological risk in four domains: deprivation, dilapidation, disconnection, and danger. <u>Deprivation</u> was measured with the Department of Community and Local Government Index of Multiple Deprivation. <u>Dilapidation</u> was measured from resident ratings of problems in their neighborhood (e.g. litter, vandalized public spaces, vacant storefronts) and independent raters’ assessments of these same problems based on the “virtual walk-through” using Google Streetview. <u>Disconnection</u> was measured from resident surveys assessing neighborhood collective efficacy and social connectedness. *Neighborhood collective efficacy* was assessed via the resident survey using a previously validated 10-item measure of social control and social cohesion ^48^. Residents were asked about the likelihood that their neighbors could be counted on to intervene in various ways if, for example: ‘children were skipping school and hanging out on a street corner,’ ‘children were spray-painting graffiti on a local building’. They were also asked how strongly they agreed that, for example: ‘people around here are willing to help their neighbors,’ ‘this is a close-knit neighborhood’ (item responses: 0-4). *Social connectedness* was assessed based on indicators of *intergenerational closure* (“If any of your neighbors’ children did anything that upset you would you feel that you could speak to their parents about it?”), *reciprocated exchange* (e.g., Would you be happy to leave your keys with a neighbor if you went away on holiday?) and *friendship ties* (e.g., Do you have any close friend that live in your neighborhood) among neighbors developed in prior research ^49^. <u>Dangerousness</u> was measured from police records of crime incidence, from neighborhood residents’ ratings of how much they feared for their safety and whether they had been victimized, and from independent raters’ assessments of neighborhood safety based on the “virtual walk-through” using Google Streetview (**Figure 5**).

**Figure 5.**
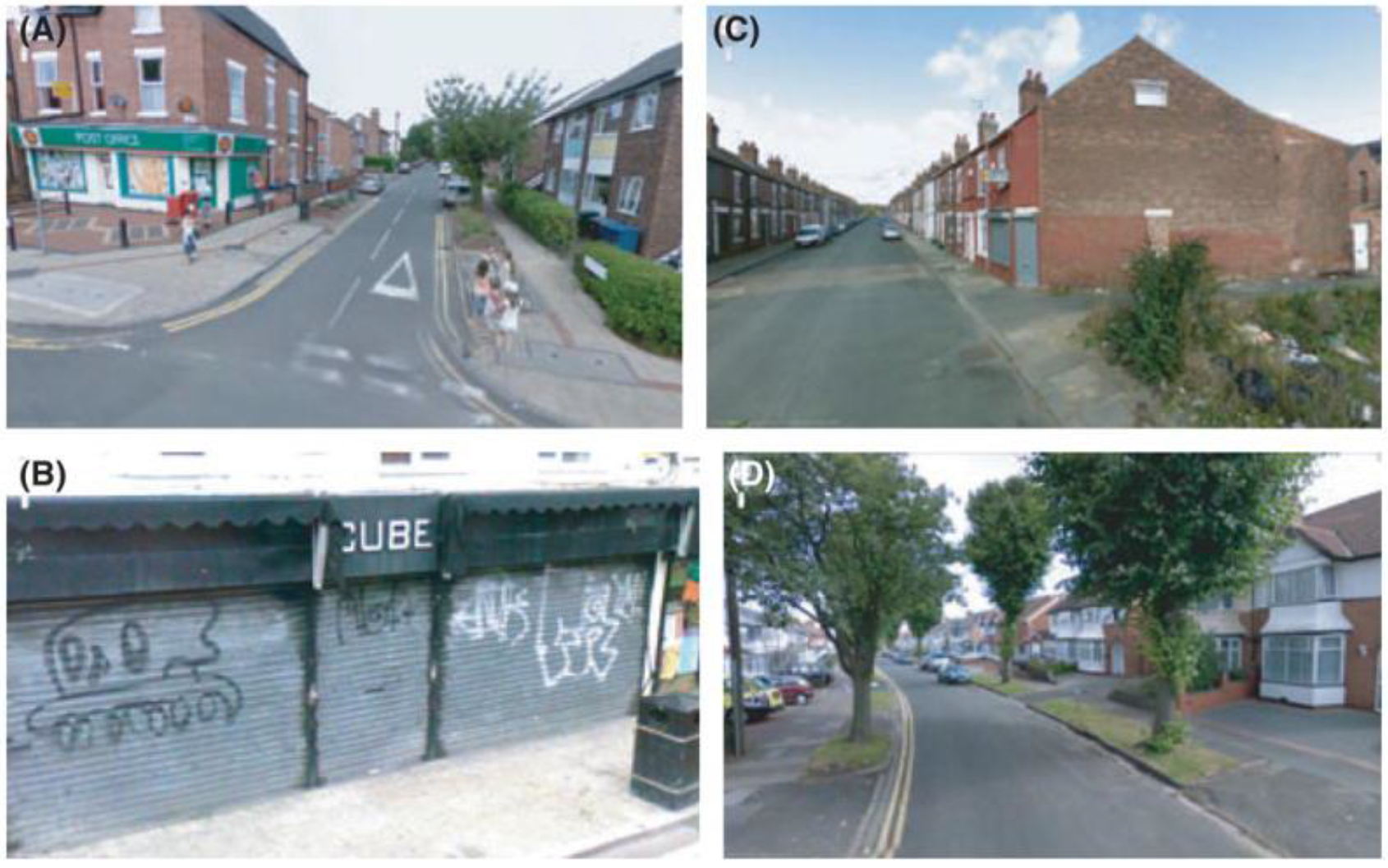
Google Street View images. (a) Well-kept neighborhood; children and amenities visible on the street; roads and sidewalks in good condition. (b) Evidence of graffitti; poorly kept sidewalk and trash container; sidewalks in fair condition. (c) Deprived residential area; vacant lot in poor condition; heavy amount of litter; sidewalks and road in poor condition. (d) Comfortably-off residential area; roads and sidewalks in good conditions; no signs of litter, graffitti or other signs of disorder

**Figure 6.**
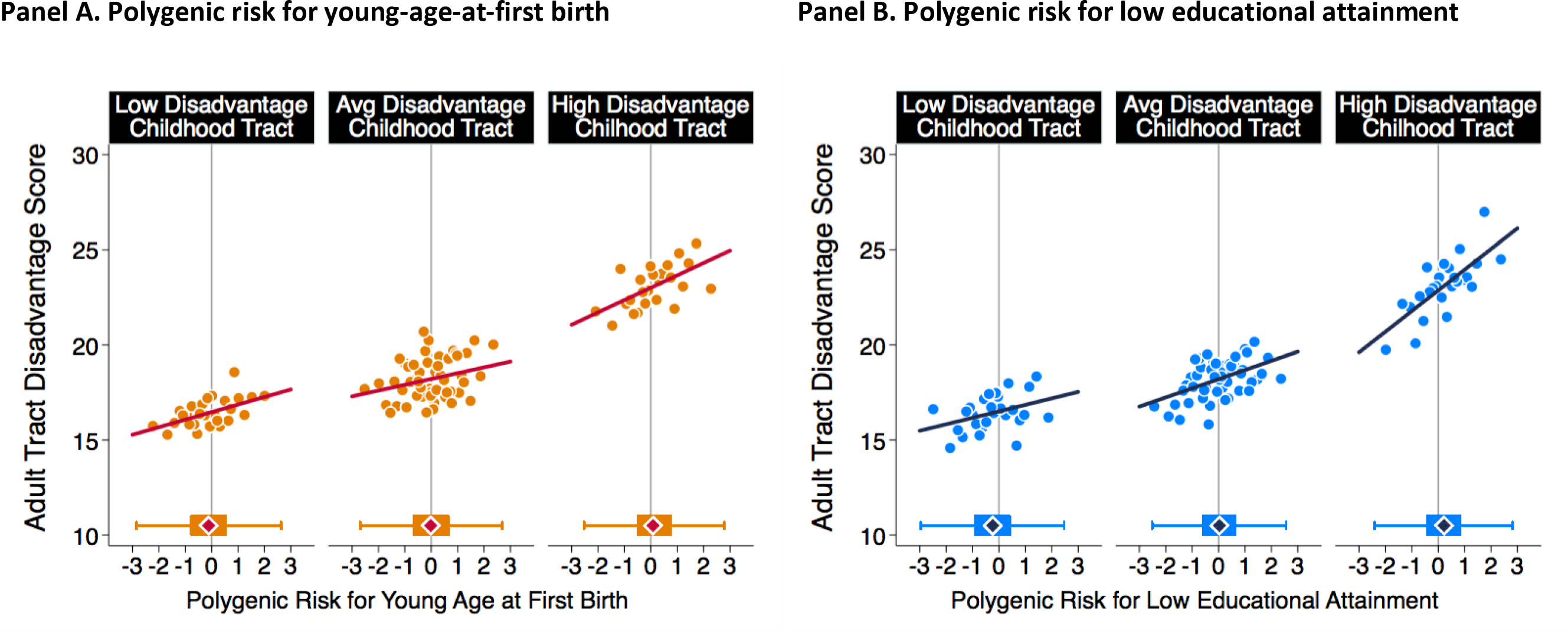
Education polygenic score association with neighborhood mobility in the Add Health Study. The figure plots polygenic risk associations with adult neighborhood disadvantage at the Census tract level for Add Health Participants who grew up in low-, middle-, and high-disadvantage Census tracts. For the figure, low-, middle-, and high-disadvantage Census tracts were defined as the bottom quartile, middle 50%, and top quartiles of the childhood tract disadvantage score distribution. The individual graphs show binned scatterplots in which each plotted point reflects average X- and Y- coordinates for a “bin” of 50 Add Health participants. The regression lines are plotted from the raw data. The box-and-whisker plots at the bottom of the graphs show the distribution of polygenic risk for each childhood-neighborhood-disadvantage category. The blue diamond in the middle of the box shows the median; the box shows the interquartile range; and the whiskers show upper and lower bounds defined by the 25^th^ percentile minus 1.5x the interquartile range and the 75^th^ percentile plus 1.5x the interquartile range, respectively. The vertical line intersecting the X-axis shows the cohort average polygenic risk. The figure illustrates three findings. First, adult participants tended to live in Census tracts with similar levels of disadvantage to the ones where they grew up. Second, children’s polygenic risks and their neighborhood disadvantage were correlated; the box plots show polygenic risk tended to be lower for participants who grew up in low-disadvantage tracts and higher for participants who grew up in high disadvantage tracts. Third, across strata of childhood neighborhood disadvantage, children at higher polygenic risk tended to move to more disadvantaged Census tracts no matter where they grew up.

For each of the four domains, we constructed a measure of ecological risk as follows. First, variables with skewed distribution were log transformed. Second, values were standardized to have M=50, SD=10. (For domains in which multiple resident survey or systematic-social-observation measures were available, we combined values within measurement method before standardizing.) Finally, scores were averaged across measurement method within each domain. The resulting scales of deprivation, dilapidation, disconnection, and danger were approximately normally distributed (**Figure 2 Panel B**). Neighborhoods’ ecological risk levels on these four measures were correlated (Pearson’s r=0.4-0.7, **Figure 2**, **Panel B**). We computed the composite Ecological-Risk Index by summing values across the four risk domains. Values were pro-rated for families with data on at least three of the four domains. Ecological-Risk Index values were available for N=1,954 children in 987 families in the genetic sample.

#### Phenotypes

We selected phenotypes for analysis that represented substantial public health and economic burden, had been linked with neighborhood risk in prior studies, were prevalent among 18-year-olds in the United Kingdom at the time data were collected, and had been subject to large-scale genome-wide association study meta-analyses: obesity, mental health problems, teen pregnancy, and poor educational outcome.

##### Obesity

Trained research workers took anthropometric measurements of study members when they were aged 18 years. BMI was computed as weight in kilograms over squared height in meters. Waist\hip ratio was calculated by dividing waist circumference by hip circumference. We defined obesity using the US Centers for Disease Control and Prevention threshold of BMI>30 and the World Health Organization recommendation of waist-hip ratio >0.90 for men and >0.85 for women ^50^. 21% of the analysis sample met at least one of these criteria, similar to prevalence for 16-24 year olds in the UK ^51^.

##### Mental Health Problems

Our measure of mental health problems is a general factor of psychopathology, the ‘p-factor,’ derived from confirmatory factor analysis of symptom-level psychopathology data collected at age 18 years, when E-Risk participants were assessed in private interviews about alcohol dependence, tobacco dependence, cannabis dependence, conduct disorder, attention-deficit hyperactivity disorder, depression, generalized anxiety disorder, post-traumatic stress disorder, eating disorder and thought/psychotic disorders ^52^. The ‘p factor’ indexes liability to develop a wide spectrum of mental-health problems ^53^. We classified E-Risk Study members reporting psychiatric symptoms one standard deviation or more above the cohort norm as having mental health problems. 17% of the analysis sample met this criterion.

##### Teen pregnancy

Getting pregnant (for women) and getting someone pregnant (for men) was assessed as part of a computer-assisted interview about reproductive behavior at the age-18 interview. 8% of the analysis sample (6% of men and 9% of women) reported a teen pregnancy.

##### Poor Educational Qualifications

Poor educational qualification was assessed by whether participants did not obtain or scored a low average grade (grade D-G) on their General Certificate of Secondary Education (GCSE). GSCEs are a standardized examination taken at the end of compulsory education at age 16 years. 23% of the analysis sample met criteria for poor educational qualifications.

##### NEET

NEET is an initialization for “Not in Education, Employment, or Training.” NEET status was assed at in-person interviews ^54^. As of the age-18 interview, 12% of Study members were NEET, similar to the UK population (as of 2010, about 14% of UK 19 year olds reported being NEET for at least one year ^55^.

### National Longitudinal Study of Adolescent to Adult Health (Add Health)

#### Sample

The National Longitudinal Study of Adolescent to Adult Health (Add Health) is an ongoing, nationally-representative longitudinal study of the social, behavioral, and biological linkages in health and developmental trajectories from early adolescence into adulthood. The cohort was drawn from a probability sample of 144 middle and high schools and is representative of American adolescents in grades 7-12 in 1994-1995. Since the start of the project, participants have been interviewed in home at four data collection waves (numbered I-IV), most recently in 2007-2008, when 15,701 Study members took part ^25^.

#### Genotyping

At the Wave IV interview in 2007-2008, saliva and capillary whole blood were collected from respondents. 15,159 of 15,701 individuals interviewed consented to genotyping, and 12,254 agreed to genetic data archiving. DNA extraction and genotyping was conducted on this archive sample using two platforms (Illumina Omni1 and Omni2.5). After quality controls, genotype data were available for 9,975 individuals. We analyzed data from the N=5,690 participants with genetically European ancestry. Imputation was conducted on SNPs “called” in more than 98% of the samples with minor allele frequency >1% using the Michigan Imputation Server (http://imputationserver.readthedocs.io/en/latest/pipeline/) and the Haplotype Reference Consortium (HRC) reference panel ^56^.

#### Polygenic Scoring

We computed polygenic scores for body-mass-index, schizophrenia and age-at-first-birth following the method described by Dudbridge ^57^ according to the procedure used in previous studies ^58^. Briefly, SNPs in the genotype database were matched to published GWAS results ^16,17^. For each of these SNPs, a loading was calculated as the number of phenotype-associated alleles multiplied by the effect-size estimated in the original GWAS. Loadings were then averaged across the SNP set to calculate the polygenic score. The Add Health Study was included in the most recent GWAS of educational attainment ^18^. We therefore obtained the polygenic score for educational attainment directly from the Social Science Genetic Association Consortium (SSGAC). SSGAC computed the score according to the methods described in the GWAS article based on a GWAS that did not include any Add Health samples.

To account for any residual population stratification within the European-descent analysis sample, we residualized polygenic scores for the first ten principal components estimated from the genome-wide SNP data ^47^ and standardized residuals to have mean=0, SD=1 to compute polygenic scores for analysis. Principal components for the Add Health European-descent sample were provided by the SSGAC.

#### Neighborhood Characteristics

We measured neighborhood-level socioeconomic disadvantage using Census-tract-level data linked to Add Health participants’ addresses when they were first interviewed in 1994-1995 and when they were most recently followed-up in 2007-8. Participants’ 1994-1995 addresses were linked with tract-level data from the 1990 Decennial Census ^59^. Participants’ addresses in 2007-2008 were linked with tract-level data from the 2005-2009 panels of the American Community Survey ^60^. For each tract, we coded proportions of female-headed households, individuals living below the poverty threshold, individuals receiving public assistance, adults with less than a high school education, and adults who were unemployed using the following system: We computed tract deciles based on the full set of tracts from which Add Health participants were sampled at Wave 1. We then scored each tract on a scale of 1-10 corresponding to the Wave 1 decile containing the tract’s value on the variable. We calculated neighborhood deprivation as the sum of decile scores across the five measures resulting in a score ranging from 0 to 50. Values were Z-transformed to have M=0 SD =1 for analysis.

Add Health analysis included all European-descent Add Health participants with available genetic and neighborhood data (N=5,325).

#### Statistical Analysis

We analyzed continuous dependent variables using linear regression models. We analyzed dichotomous dependent variables using Poisson regression models to estimate risk ratios (RR). In models testing polygenic and neighborhood risks for health and social problems, health and social problems were specified as the dependent variables and polygenic and neighborhood risks were specified as predictor variables. We tested statistical independence of polygenic risk information from family history risk information using multivariate regression with family history measures included as covariates alongside polygenic scores. In models testing for association between polygenic and neighborhood risks, polygenic scores were specified as dependent variables and neighborhood risks were specified as predictor variables. We tested if associations between children’s neighborhood risks and polygenic risks were correlated because both were inherited from their parents using multivariate regression with mother’s polygenic scores included as covariates alongside neighborhood risk measures. We tested polygenic risk associations with neighborhood mobility using the mobility model from previous work ^61^; adult neighborhood disadvantage score was regressed on the participant’s polygenic score, their child neighborhood disadvantage score, and covariates. For all models, we accounted for non-independence of observations of siblings within families by clustering standard errors at the family level. For models testing polygenic score associations with neighborhood conditions in the E-Risk data, only one member of each monozygotic twin pair was included in analysis. (For these models, monozygotic twins would have identical values for predictors and outcomes.) All models were adjusted for sex. Add Health models were adjusted for year of birth. (Year of birth did not vary in the E-Risk cohort.)

We conducted post-hoc power analyses to provide context for interpretation of the associations we observed. We conducted power analysis using the “power” command in the Stata software^62^. Both the E-Risk and Add Health samples had >80% power to detect associations with effect-size r=0.1 in all analyses. Power analysis for tests of polygenic score associations with neighborhood risk is shown in **Supplementary Figure 3**.

## Acknowledgement

The E-Risk Study is funded by the Medical Research Council (UKMRC grant G1002190). Additional support was provided by NICHD grant HD077482, Google, and by the Jacobs Foundation. The Add Health Study is supported by Eunice Kennedy Shriver National Institute of Child Health and Human Development grants P01HD31921, R01HD073342, and R01HD060726, with cooperative funding from 23 other federal agencies and foundations. DWB & CLO were supported by fellowships from Jacobs Foundation. CLO is supported by the Canadian Institute for Advanced Research. BWD is supported by Russell Sage Foundation award 961704. We are grateful to the E-Risk Study mothers and fathers, the twins, and the twins’ teachers and the Add Health Study participants and their parents for their participation. Our thanks to CACI, Google Streetview, and to members of the E-Risk team for their dedication, hard work, and insights.

## Supplementary Methods

### Ecological-Risk Assessment Measures of Neighborhood Disadvantage

Ecological risk assessment was conducted by combining information from four independent sources of data: Geodemographic data from local governments, official crime data from the UK police, Google Streetview-based Systematic Social Observation (SSO), and surveys of neighborhood residents.

#### 1. Geodemographic Data from Local Governments

We obtained information about the Index of Multiple Deprivation from the Department for Communities and Local Government. The Index is the official measure of relative deprivation for neighborhoods in England. Every small area in England is ranked from 1 (most deprived area) to 32,844 (least deprived area), these rankings are then converted into deciles. The Index of Multiple Deprivation is created based on 37 separate indicators, that are organized in seven domains of deprivation (Employment Deprivation; Health Deprivation and Disability; Education, Skills and Training Deprivation; Crime; Barriers to Housing and Services; and Living Environment Deprivation), and combined with appropriate weights to calculate the Index of Multiple Deprivation (IMD). Households were assigned a neighborhood IMD based on street address at the time of the age-5, age-7, age-10, and age-12 in-home visits. We analyzed the average IMD value across these four measurements.

#### 2. Official Crime Data

We measured local area crime by mapping a 1-mile radius around each E-Risk Study family’s home and tallying the total number of crimes that occurred in the area each month. Street-level crime data, including information on the type of crime, date of occurrence, and approximate location, were accessed online as part of an open data sharing effort about crime and policing in England and Wales (https://data.police.uk/) and geocoded to the home address of the study members. An Application Program Interface was used to extract street-level crime data for each of the geospatial coordinates marking the family’s home. For a full description see: https://data.police.uk/about/#location-anonymisation.

#### 3. Google Street View Virtual Systematic Social Observation (SSO)

The Google Streetview-SSO consisted of trained raters taking a “virtual walk” through the neighborhoods of the E-Risk families. Raters then coded neighborhoods based on what they saw on that virtual walk. Street View is a freely available tool that generates panoramic street-level views using high definition images taken from camera-equipped cars. Signals from global positioning devices are used to accurately position images in the online maps. To avoid gaps in the imagery, adjacent cameras on the car take overlapping pictures and the images are then stitched together to create a continuous 360-degree image of the street. Images are then smoothed and re-projected onto a sphere to create the image displayed in Street View (see **Figure 5**). To protect the privacy of individuals, face- and license-blurring technology is applied to ensure that people on the street and cars in the photographs cannot be identified. Google Street View came online in the United Kingdom in March 2009 and by March 2010, 94% of the E-risk children’s neighborhoods were available for viewing. The Google Street View Systematic Social Observation (SSO) was completed by adapting SSO instruments for the virtual context and training raters to reliably code neighborhood features while taking a virtual walk down the street. We have reported full details of the Google Street View SSO method, inter-rater reliability and predictive validity of the measures elsewhere (21). We analyzed Google Street View SSO measures of environmental decay and disorder and perceived dangerousness.

#### 4. Resident Surveys

A survey of residents living alongside E-Risk families was conducted when the children were 13-14 years of age to capture neighborhood-level social processes that cannot easily be captured via official records or direct observation.

The sampling frame for the Neighborhood Survey was drawn using UK-Info Pro V13 http://www.192.com/products/. The survey responses were anonymous; no identifying information was collected. In Britain, a postcode area typically contains 15 households, with at most 100 households (e.g., a large apartment block). Therefore, survey respondents were typically living on the same street or within the same apartment block as the children in our study. Surveys were mailed to every household in the postcode registered to the electoral role, with the exception of the E-Risk family, resulting in 20,529 surveys being mailed to households to capture information on E-Risk families. On average, we received 5 (SD=3) completed surveys per neighborhood (range= 0-18 respondents). We achieved at least 3 responses for 80% of target neighborhood and at least 2 responses for 95% (resulting in a total of 5601 completed questionnaires). Survey responses were received for N=1,077 of the 1,116 families in the study. We analyzed survey measures of the following neighborhood-level social processes: fear of crime, direct victimization, neighborhood problems, and social disconnectedness.

### Calculations to Evaluate How Much of the Neighborhood Gradient in Risk for Poor Educational Attainment and NEET Status Might Be Explained by Gene-Neighborhood Correlation Between the Education Polygenic Score and ACORN and Ecological-Risk Score Measures of Neighborhood Risk

We evaluated the extent to which gene-neighborhood correlations between GWAS discoveries for educational attainment and measures of neighborhood risk might account for the neighborhood gradient in risk for poor educational outcomes and NEET status.

First, we computed the average polygenic risk for E-Risk participants living in very low-and very high-risk neighborhoods (ACORN scores of 1 and 4; Ecological Risk Scores of 150 and 275) based on the regressions reported in **Supplemental Table S4** (see also **Figure 4** of the main text). For the very low- and very high-risk neighborhoods, the predicted values of the education polygenic score were −0.30 and 0.23 for ACORN and −0.21 and 0.31 for the Ecological-Risk Score, a difference of about 0.5 standard deviations. (The education polygenic score is reversed in our analysis relative to the original GWAS, so high values correspond to a genetic prediction of low educational attainment).

Next, we computed predicted values of the phenotypes for participants with those polygenic scores based on the regression reported in **Supplemental Table S1**, **Panel A** (see also **Figure 1** of the main text). The predicted proportions of individuals with poor educational qualifications corresponding to the low and high polygenic risk values were 19% and 24% based on the ACORN predictions and 20% and 24% based on the Ecological Risk Score predictions, a difference of 4-5%. The predicted proportions of individuals who were NEET corresponding to the low and high polygenic risk values were 11% and 12% based on the ACORN predictions and 11% and 13% based on the Ecological Risk Score predictions, a difference of 1-2%.

Finally, we computed predicted values of the phenotypes for participants who grew up in very low- and very high-risk neighborhoods based on the regressions reported in **Supplemental Table S3** (see also **Figure 2** of the main text). The predicted proportions of individuals with poor educational qualifications corresponding to the very low- and very high-risk neighborhoods were 8% and 38% based on the ACORN predictions and 14% and 46% based on the Ecological Risk Score predictions, a difference of 30-32%. The predicted proportions of individuals who were NEET corresponding to the very low- and very high-risk neighborhoods were 7% and 20% based on the ACORN predictions and 6% and 27% based on the Ecological Risk Score predictions, a difference of 13-21%.

To summarize, we observed a difference in polygenic risk for low educational attainment between very low-risk and very high-risk neighborhoods of roughly one half of one standard deviation. Based on our analysis, this difference in genetic risk could account for a difference in the prevalence of poor educational qualifications between very low-risk and very high-risk neighborhoods of 4-5%. The observed difference in the prevalence of poor educational qualifications between very low-risk and very high-risk neighborhoods was 30-32%, 6-7 times larger. The pattern of results was similar for NEET status. Based on our analysis, genetic risk could account for a neighborhood difference of 1-2% in prevalence. The observed difference in prevalence was 13-21%, an order of magnitude larger.

**Supplemental Table S1.** Polygenic score and family history associations with E-Risk children’s health and social problems. **Panel A** shows effect-sizes for polygenic risk associations with children’s health and social problems. Effect-sizes are relative risks (RR) estimated from Poisson regression models for a 1 SD increase in polygenic risk. Models included all E-Risk Study members with available genotype and phenotype data (N=1,837 for obesity; N=1,825 for teen pregnancy; N=1,863 for mental health problems; N=1,860 for poor educational qualifications; N=1,863 for NEET status). All models were adjusted for sex. Nesting of twins within families was accounted for by clustering standard errors at the family level.

| Phenotype | Polygenic Score | RR | 95% CI | p-value |
| --- | --- | --- | --- | --- |
| Obesity | Body-mass Index | 1.26 | [1.14-1.38] | 1.99E-06 |
| Mental Health Problems | Schizophrenia | 1.13 | [1.02-1.26] | 0.022 |
| Teen Pregnancy | Age-at-first-birth (reversed) | 1.40 | [1.19-1.64] | 3.24E-05 |
| Poor Educational Qualification | Educational | 1.47 | [1.34-1.60] | 8.96E-18 |
| NEET | Attainment (reversed) | 1.32 | [1.15-1.52] | 7.42E-05 |

| Phenotype | Polygenic Score | r | 95% CI | p-value |
| --- | --- | --- | --- | --- |
| Body-mass Index | Body-mass Index | 0.26 | [0.21-0.31] | 1.04E-21 |
| Mental Health Problems | Schizophrenia | 0.06 | [0.01-0.11] | 2.30E-02 |
| Educational Attainment | Educational Attainment (reversed) | 0.28 | [0.23-0.33] | 1.04E-25 |

**Panel B** shows correlations between measures of polygenic and family history risk. Maternal body-mass-index (BMI) was measured from mother’s self-reported height and weight when E-Risk participants were aged 12 years (N=900 mothers of 1,780 participants; M=26, SD=6); family history of psychiatric hospitalization was measured based on family histories collected during interviews with children’s mothers as the proportion of relatives with a hospitalization (N=970 mothers of 1,920 participants; M=0.07, SD=0.13); mother’s age at first birth was collected as part of screening for enrollment in the study (N=1,011 mothers of 1,999 participants; M=24 years, SD=6); the highest education of either parent was collected during interviews when participants were aged 5 years (N=1,011 families of 1,999 participants; 12% held no educational credentials, 12% held GCSE Level-1 credentials; 35% held GCSE Level-2 credentials; 41% held GCSE Level-3 or higher credentials).

| **Polygenic Risk Measure** | **Family History Measure** | **Pearson r** |
| --- | --- | --- |
| BMI Polygenic Score | Maternal BMI | 0.13 |
| Schizophrenia Polygenic Score | Family History of Psychiatric Hospitalization | 0.01 |
| Age-at-first-birth Polygenic Score | Mother's age at first birth | 0.16 |
| Educational Attainment Polygenic Score | Parents' Education | 0.23 |

**Panel C** shows effect-sizes for polygenic risk and family-history associations with children’s health and social problems. Samples are restricted to children with available phenotype, genotype, and family-history information (N=1,666 for obesity; N=1,812 for mental health problems; N=1,825 for teen pregnancy; N=1,860 for poor educational qualifications and NEET status). The first column (M1) reports the effect-size for polygenic risk. The second column (M2) reports the effect-size for family-history risk. The third column (M3) reports the multivariate-adjusted effect-sizes for polygenic risk and family-history risk from a model that includes both risk factors. All models were adjusted for sex. Nesting of twins within families was accounted for by clustering standard errors at the family level.

|  | **M1** | **M2**<br><u>RR [95% CI]</u> | **M3** |
| --- | --- | --- | --- |
| **Obesity** |  |  |  |
| BMI Polygenic Score | 1.19<br>[1.08-1.32] |  | 1.17<br>[1.03-1.33] |
| Family History<br>(maternal BMI) |  | 1.24<br>[1.14-1.34] | 1.21<br>[1.11-1.32] |
| **Mental Health Problems** |  |  |  |
| Schizophrenia Polygenic<br>Score | 1.13<br>[1.01-1.26] |  | 1.04<br>[0.91-1.2] |
| Family History<br>(maternal family history of |  | 1.19<br>[1.08-1.3] | 1.18<br>[1.07-1.3] |
| **Teenaged Pregnancy** |  |  |  |
| Age-at-first-birth Polygenic<br>Score (reversed) | 1.40<br>[1.19-1.64] |  | 1.18<br>[0.97-1.43] |
| Family History<br>(mother's age at first birth) |  | 2.34<br>[1.80-3.03] | 2.21<br>[1.65-2.97] |
| **Poor Educational Qualifications** |  |  |  |
| Educational Attainment<br>Polygenic Score (reversed) | 1.47<br>[1.34-1.60] |  | 1.19<br>[1.07-1.33] |
| Family History<br>(parents' education) |  | 1.69<br>[1.57-1.82] | 1.57<br>[1.44-1.71] |
| **NEET** |  |  |  |
| Educational Attainment<br>Polygenic Score (reversed) | 1.32<br>[1.15-1.52] |  | 1.11<br>[0.95-1.31] |
| Family History (parents'<br>education) |  | 1.72<br>[1.53-1.94] | 1.62<br>[1.41-1.87] |

**Supplemental Table S2.** Means and standard deviations of neighborhood measures and their correlations with one another.

|  |  |  | **Pearson Correlation** |  |  |  |  |  |
| --- | --- | --- | --- | --- | --- | --- | --- | --- |
|  | M | SD | (1) | (2) | (3) | (4) | (5) | (6) |
| (1) ACORN | 2.49 | 1.10 |  |  |  |  |  |  |
| (2) Composite Ecological-Risk Index | 198.46 | 32.77 | 0.65 |  |  |  |  |  |
| (3) Deprived | 49.54 | 9.85 | 0.53 | 0.76 |  |  |  |  |
| (4) Dilapidated | 49.73 | 9.79 | 0.52 | 0.85 | 0.50 |  |  |  |
| (5) Disconnected | 49.72 | 9.94 | 0.50 | 0.81 | 0.44 | 0.58 |  |  |
| (6) Dangerous | 49.39 | 9.88 | 0.61 | 0.89 | 0.58 | 0.72 | 0.64 |  |

**Supplemental Table S3.**
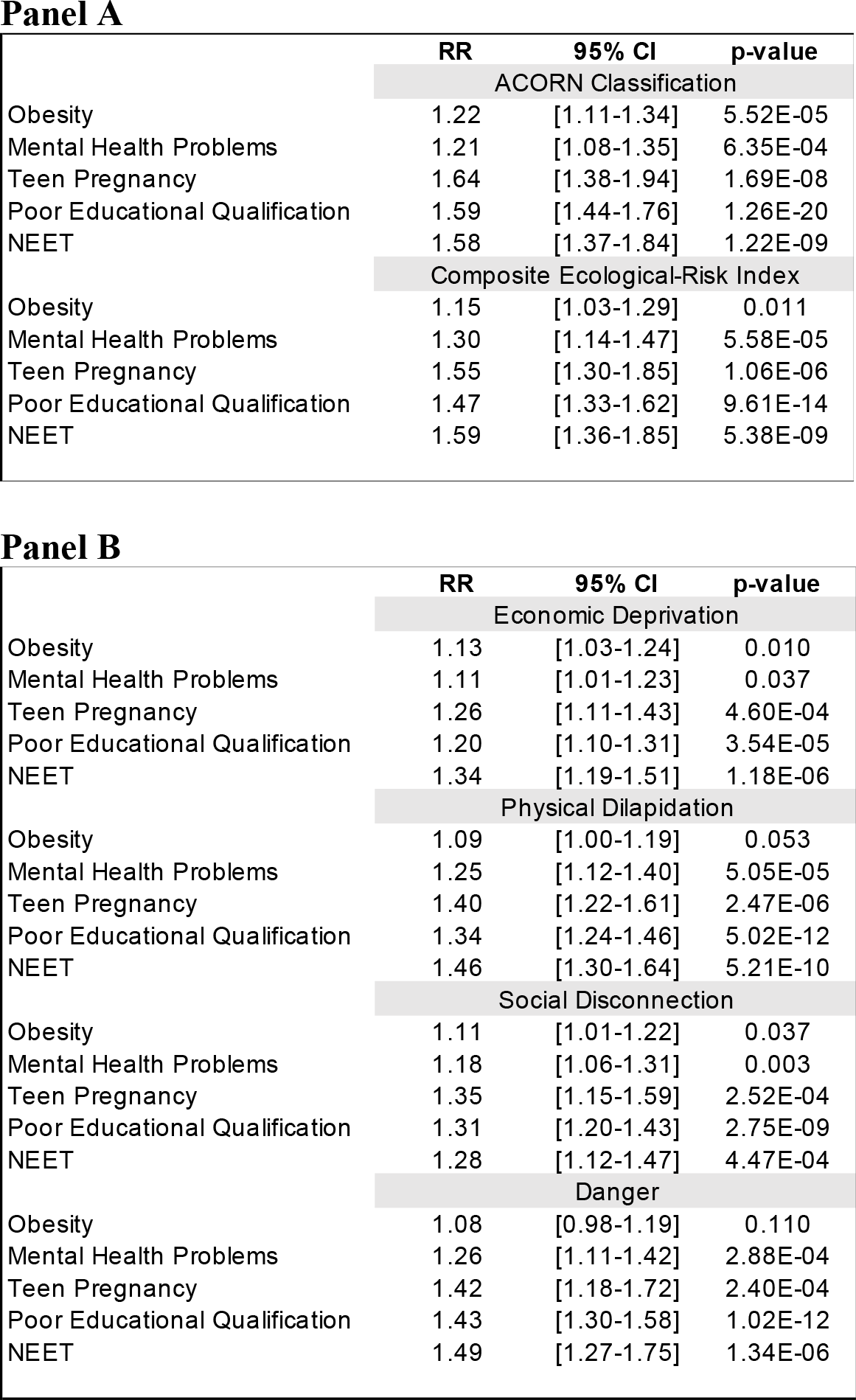
Associations between neighborhood disadvantage measures and children’s health and social problems. Table shows relative risks (RR) estimated from Poisson regression models. Nesting of twins within families was accounted for by clustering standard errors at the family level. Panel A shows results for ACORN (N=1,857) and the composite Ecological-Risk Index (N=1,822). Effect-sizes for ACORN classification are reported for a 1-category increase in social disadvantage. Effect-sizes for composite Ecological-Risk Index are reported for a 1-SD increase in ecological risk. Panel B shows results for individual ecological-risk measures. Effect-sizes are reported for a 1-SD increase in ecological risk.

Panel A
|  | RR | 95% CI | p-value |
| --- | --- | --- | --- |
| ACORN Classification |  |  |  |
| Obesity | 1.22 | [1.11-1.34] | 5.52E-05 |
| Mental Health Problems | 1.21 | [1.08-1.35] | 6.35E-04 |
| Teen Pregnancy | 1.64 | [1.38-1.94] | 1.69E-08 |
| Poor Educational Qualification | 1.59 | [1.44-1.76] | 1.26E-20 |
| NEET | 1.58 | [1.37-1.84] | 1.22E-09 |
| Composite Ecological-Risk Index |  |  |  |
| Obesity | 1.15 | [1.03-1.29] | 0.011 |
| Mental Health Problems | 1.30 | [1.14-1.47] | 5.58E-05 |
| Teen Pregnancy | 1.55 | [1.30-1.85] | 1.06E-06 |
| Poor Educational Qualification | 1.47 | [1.33-1.62] | 9.61E-14 |
| NEET | 1.59 | [1.36-1.85] | 5.38E-09 |

Panel B
|  | RR | 95% CI | p-value |
| --- | --- | --- | --- |
| Economic Deprivation |  |  |  |
| Obesity | 1.13 | [1.03-1.24] | 0.010 |
| Mental Health Problems | 1.11 | [1.01-1.23] | 0.037 |
| Teen Pregnancy | 1.26 | [1.11-1.43] | 4.60E-04 |
| Poor Educational Qualification | 1.20 | [1.10-1.31] | 3.54E-05 |
| NEET | 1.34 | [1.19-1.51] | 1.18E-06 |
| Physical Dilapidation |  |  |  |
| Obesity | 1.09 | [1.00-1.19] | 0.053 |
| Mental Health Problems | 1.25 | [1.12-1.40] | 5.05E-05 |
| Teen Pregnancy | 1.40 | [1.22-1.61] | 2.47E-06 |
| Poor Educational Qualification | 1.34 | [1.24-1.46] | 5.02E-12 |
| NEET | 1.46 | [1.30-1.64] | 5.21E-10 |
| Social Disconnection |  |  |  |
| Obesity | 1.11 | [1.01-1.22] | 0.037 |
| Mental Health Problems | 1.18 | [1.06-1.31] | 0.003 |
| Teen Pregnancy | 1.35 | [1.15-1.59] | 2.52E-04 |
| Poor Educational Qualification | 1.31 | [1.20-1.43] | 2.75E-09 |
| NEET | 1.28 | [1.12-1.47] | 4.47E-04 |
| Danger |  |  |  |
| Obesity | 1.08 | [0.98-1.19] | 0.110 |
| Mental Health Problems | 1.26 | [1.11-1.42] | 2.88E-04 |
| Teen Pregnancy | 1.42 | [1.18-1.72] | 2.40E-04 |
| Poor Educational Qualification | 1.43 | [1.30-1.58] | 1.02E-12 |
| NEET | 1.49 | [1.27-1.75] | 1.34E-06 |

**Supplemental Table S4.**
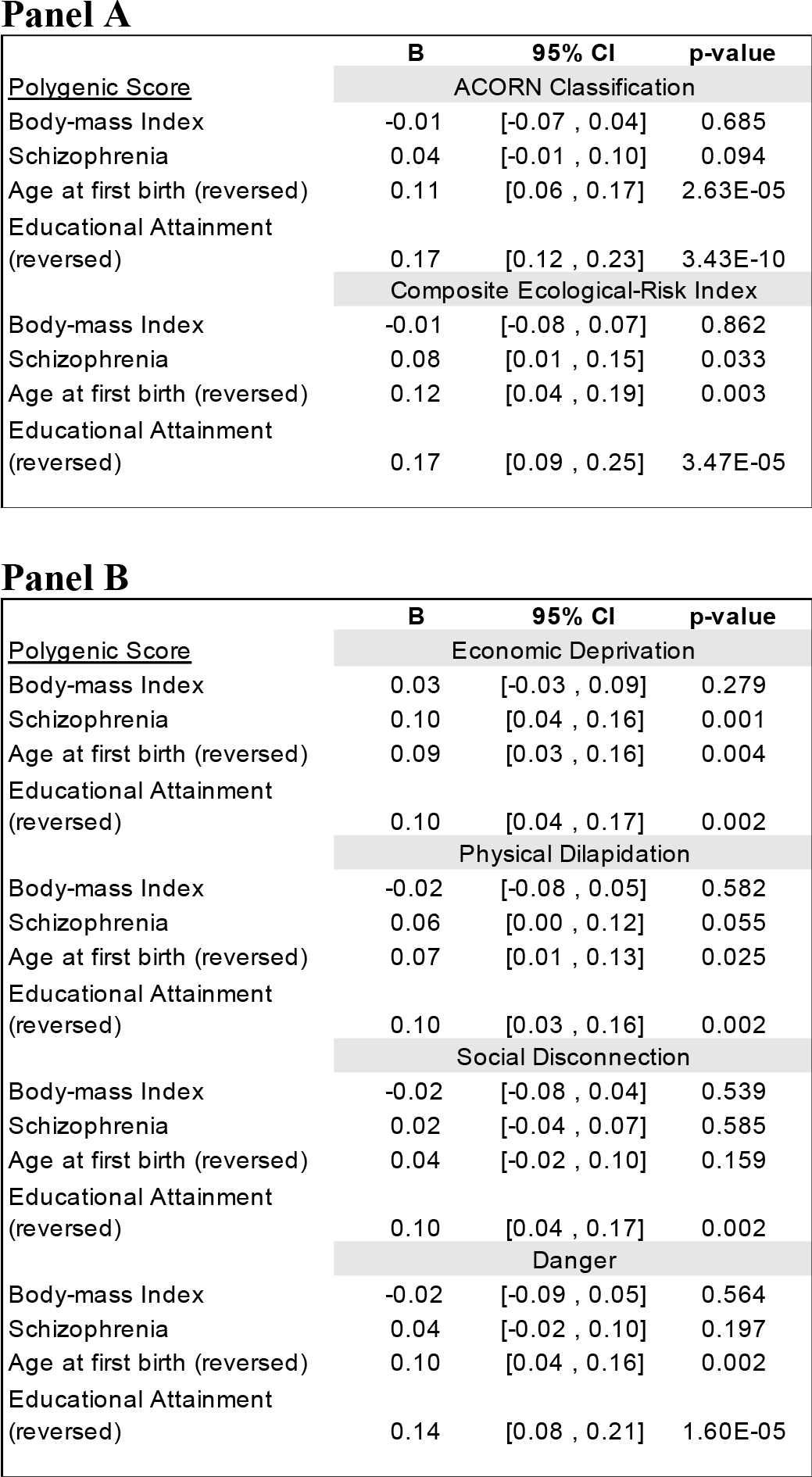
Associations between neighborhood disadvantage and children’s polygenic risk for obesity, schizophrenia, young age at first birth, and low educational attainment. Table shows unstandardized coefficients from linear regression models. Nesting of twins within families was accounted for by clustering standard errors at the family level. Only one monozygotic twin from each pair was included in analysis because monozygotic twins share identical neighborhood disadvantage and polygenic score values. Panel A shows results for ACORN (N=1,439) and the composite Ecological-Risk Index (N=1,412). Coefficients can be interpreted as expected SD increase in polygenic risk per unit increase in ACORN disadvantage classification and SD increase in polygenic score per SD increase in composite Ecological-Risk Index. Panel B shows results for individual ecological-risk measures. Coefficients can be interpreted as expected SD increase in polygenic risk per SD increase in ecological-risk.

**Supplemental Table S5.** Association between neighborhood disadvantage and mother’s polygenic risk for obesity, schizophrenia, young age at first birth, and low educational attainment. Table shows unstandardized coefficients from linear regression models. Panel A shows results for ACORN (N=838) and the composite Ecological-Risk Index (N=823). Coefficients can be interpreted as expected SD increase in polygenic risk per unit increase in ACORN disadvantage classification and SD increase in polygenic score per SD increase in composite Ecological-Risk Index. Panel B shows results for individual ecological-risk measures. Coefficients can be interpreted as expected SD increase in polygenic risk per SD increase in ecological-risk.

Panel A
|  | **B** | **95% CI** | **p-value** |
| --- | --- | --- | --- |
| <u>Polygenic Score</u> | ACORN Classification |  |  |
| Body-mass Index | 0.00 | [-0.07 , 0.06] | 0.881 |
| Schizophrenia | 0.03 | [-0.03 , 0.09] | 0.381 |
| Age at first birth (reversed) | 0.13 | [0.07 , 0.19] | 4.89E-05 |
| Educational Attainment (reversed) | 0.18 | [0.12 , 0.25] | 5.76E-09 |
|  | Composite Ecological-Risk Index |  |  |
| Body-mass Index | 0.03 | [-0.06 , 0.11] | 0.519 |
| Schizophrenia | 0.02 | [-0.07 , 0.10] | 0.713 |
| Age at first birth (reversed) | 0.18 | [0.10 , 0.26] | 2.17E-05 |
| Educational Attainment (reversed) | 0.17 | [0.09 , 0.26] | 5.31E-05 |

Panel B
|  | **B** | **95% CI** | **p-value** |
| --- | --- | --- | --- |
| <u>Polygenic Score</u> | Economic Deprivation |  |  |
| Body-mass Index | 0.03 | [-0.04 , 0.09] | 0.408 |
| Schizophrenia | 0.03 | [-0.04 , 0.10] | 0.383 |
| Age at first birth (reversed) | 0.11 | [0.05 , 0.18] | 0.001 |
| Educational Attainment (reversed) | 0.14 | [0.07 , 0.20] | 5.84E-05 |
|  | Physical Dilapidation |  |  |
| Body-mass Index | 0.02 | [-0.05 , 0.09] | 0.550 |
| Schizophrenia | 0.01 | [-0.06 , 0.09] | 0.710 |
| Age at first birth (reversed) | 0.14 | [0.07 , 0.21] | 5.05E-05 |
| Educational Attainment (reversed) | 0.08 | [0.01 , 0.15] | 0.025 |
|  | Social Disconnection |  |  |
| Body-mass Index | 0.02 | [-0.05 , 0.08] | 0.620 |
| Schizophrenia | -0.02 | [-0.09 , 0.05] | 0.493 |
| Age at first birth (reversed) | 0.09 | [0.02 , 0.16] | 0.012 |
| Educational Attainment (reversed) | 0.12 | [0.05 , 0.19] | 1.15E-03 |
|  | Danger |  |  |
| Body-mass Index | 0.02 | [-0.06 , 0.09] | 0.658 |
| Schizophrenia | 0.01 | [-0.06 , 0.08] | 0.686 |
| Age at first birth (reversed) | 0.14 | [0.07 , 0.21] | 1.25E-04 |
| Educational Attainment (reversed) | 0.14 | [0.06 , 0.21] | 1.94E-04 |

**Supplemental Table S6.** Effect-sizes for associations between children’s neighborhood and polygenic risks before and after covariate adjustment for their mothers’ polygenic risk. Table shows unstandardized coefficients from linear regression model of associations between neighborhood disadvantage and polygenic risk (rGE). Unadjusted associations are parallel to results reported in Supplemental Table 4 with a sample restricted to the subset of E-Risk participants for whom maternal polygenic score values were available (N=1,205 participants in 838 families for ACORN analysis and N=1,185 participants in 823 families for Ecological-Risk Index analysis). Standard errors were clustered at the family level to account for non-independence of data. Only one monozygotic twin from each pair was included in analysis because monozygotic twins share identical neighborhood disadvantage and polygenic score values.

|  | ACORN |  | Ecological-Risk Index |  |
| --- | --- | --- | --- | --- |
|  | B | [95% CI] | B | [95% CI] |
| BMI Polygenic Score |  |  |  |  |
| Unadjusted rGE | -0.01 | [-0.08-0.05] | -0.01 | [-0.09-0.08] |
| Maternal PGS-adjusted | -0.01 | [-0.06-0.04] | -0.02 | [-0.09-0.05] |
| Schizophrenia Polygenic Score |  |  |  |  |
| Unadjusted rGE | 0.04 | [-0.02-0.10] | 0.05 | [-0.03-0.13] |
| Maternal PGS-adjusted | 0.03 | [-0.01-0.08] | 0.05 | [-0.02-0.11] |
| Age-at-first-birth Polygenic Score (reversed) |  |  |  |  |
| Unadjusted rGE | 0.11 | [0.05-0.17] | 0.12 | [0.04-0.21] |
| Maternal PGS-adjusted | 0.04 | [-0.01-0.09] | 0.02 | [-0.04-0.09] |
| Educational Attainment Polygenic Score (reversed) |  |  |  |  |
| Unadjusted rGE | 0.16 | [0.11-0.22] | 0.15 | [0.07-0.24] |
| Maternal PGS-adjusted | 0.07 | [0.02-0.12] | 0.06 | [-0.01-0.13] |

**Supplemental Figure S1.**
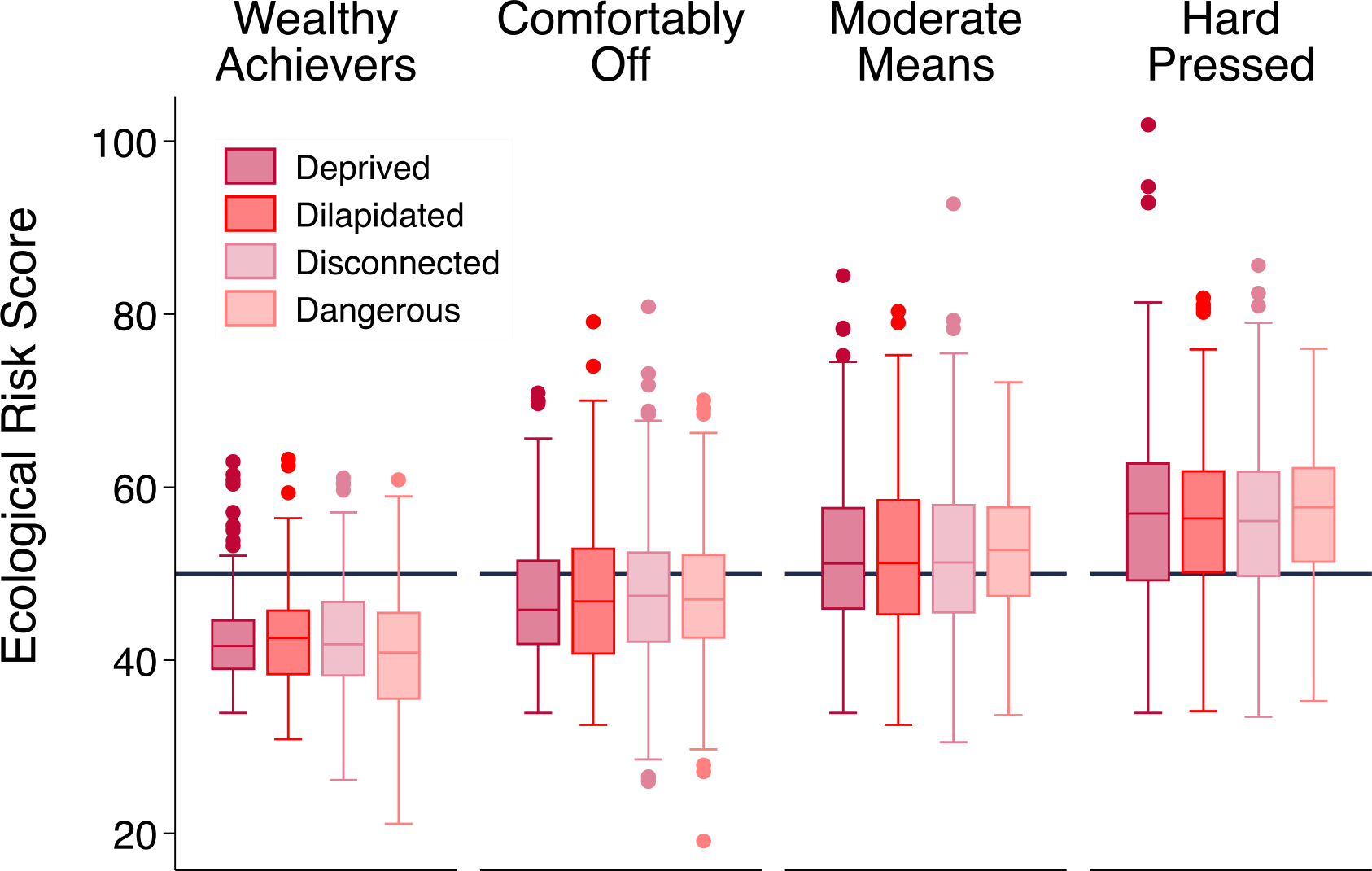
Distributions of ecological-risk assessments within ACORN neighborhood classifications. We used two different methods to measure neighborhood disadvantage: A geodemographic index (ACORN) and an Ecological Risk Index based on surveys of neighborhood residents, electronic record data, and Systematic Social Observation using Google Streetview. Neighborhoods with more socially disadvantaged ACORN classification were also more disadvantaged as measured by ecological-risk assessment. The Figure graphs distributions of each ecological-risk measure within each ACORN classification for N=1,008 families for which both ACORN and ecological-risk assessment measurements were available. For each ecological-risk measure, average risk trended upwards from the least disadvantaged ACORN classification (Wealthy Achievers) to the most disadvantaged ACORN classification (Hard Pressed).

**Supplemental Figure S2.**
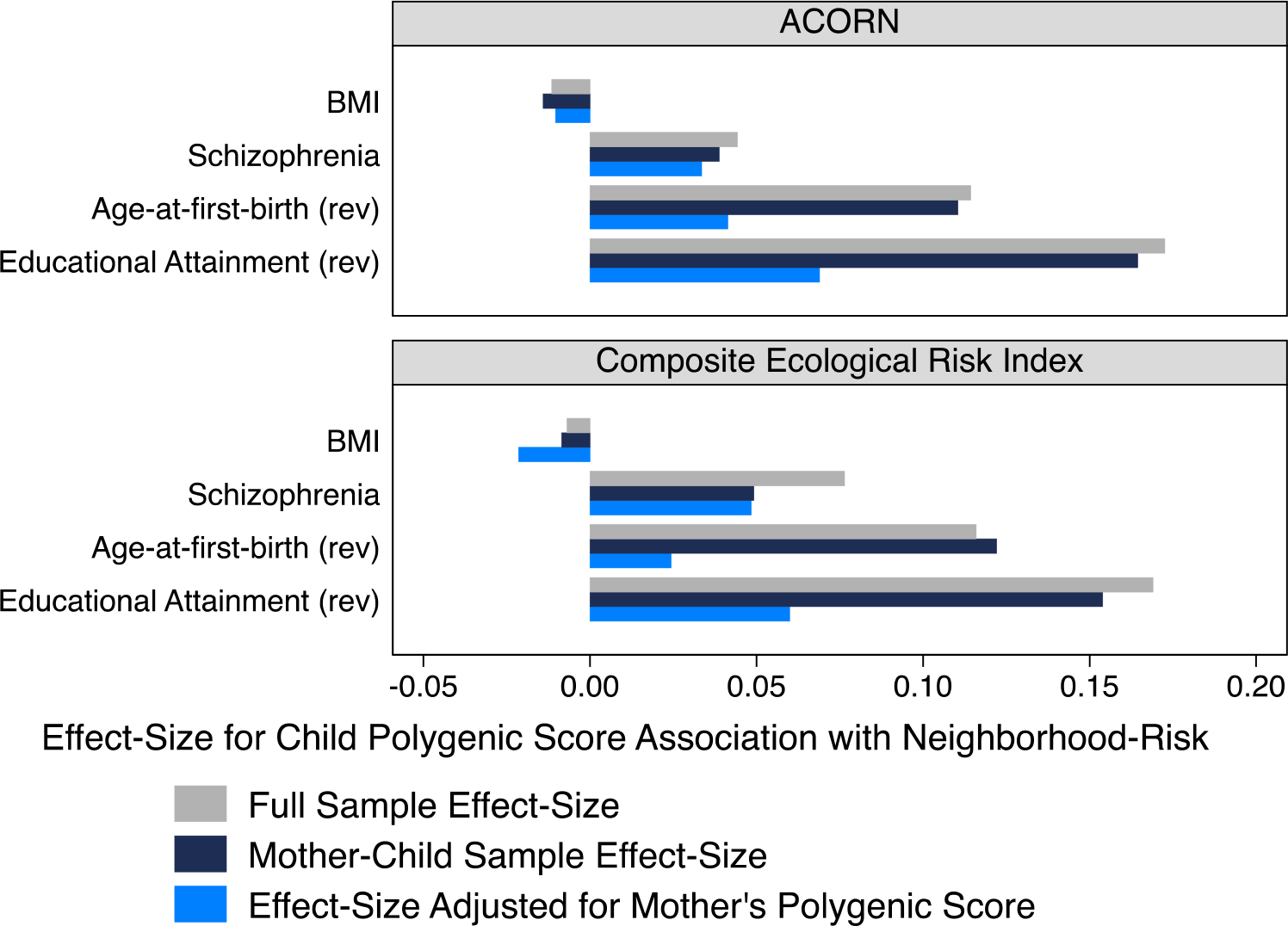
Effect-sizes for associations between children’s neighborhood and polygenic risks before and after covariate adjustment for their mothers’ polygenic risk. The figure shows effect-sizes for associations between children’s polygenic and neighborhood risks in the full E-Risk sample (gray bars, N=1,439 for ACORN, 1,412 for Ecological-Risk Index, see also Supplemental Table 4), in the mother-child sub-sample which included families with genetic data on mothers and children (dark blue bars, N=1,205 for ACORN, 1,185 for the Ecological-Risk Index), and in the mother-child sub-sample after covariate adjustment for mother’s polygenic score (light blue bars). Nesting of twins within families was accounted for by clustering standard errors at the family level. The top panel shows results for ACORN neighborhood disadvantage classification. Coefficients can be interpreted as expected SD increase in polygenic risk per unit increase in ACORN disadvantage classification. The bottom panel shows results for the composite Ecological-Risk Index. Coefficients can be interpreted as expected SD increase in polygenic risk per SD increase in ecological-risk.

**Supplemental Figure S3.**
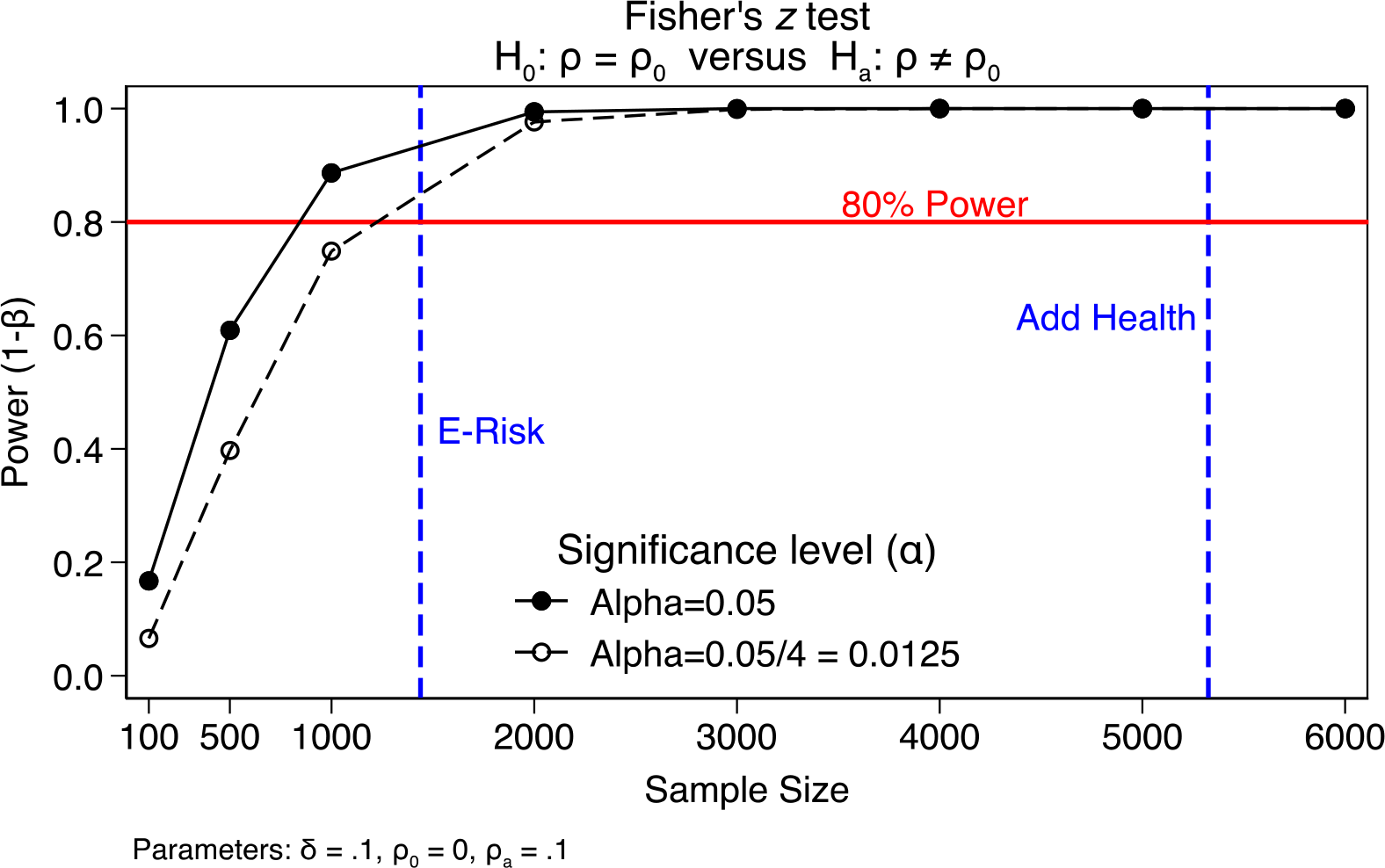
Power for testing genetic associations with neighborhood risk. The figure plots statistical power on the y-axis against sample-size on the x-axis for testing effect-sizes of r=0.1 against a null hypothesis of r=0. Power is plotted for the conventional alpha threshold of 0.05 as well as an alpha threshold corrected for testing 4 polygenic scores (0.05/4=0.0125). The sample-sizes for E-Risk and Add Health tests of genetic association with neighborhood risk are denoted by vertical blue lines. The threshold of 80% power is denoted with a horizontal red line. The graph shows that both samples have >80% power to test associations with effect-size of r=0.1.

